# The asymmetric *Pitx2* regulates intestinal muscular-lacteal development and protects against fatty liver disease

**DOI:** 10.1101/2021.06.11.447753

**Authors:** Shing Hu, Aparna Mahadevan, Isaac F. Elysee, Joseph Choi, Nathan R. Souchet, Gloria H. Bae, Alessandra K. Taboada, Gerald E. Duhamel, Carolyn S. Sevier, Ge Tao, Natasza A. Kurpios

## Abstract

Intestinal lacteals are the essential lymphatic channels for absorption and transport of dietary lipids and drive pathogenesis of debilitating metabolic diseases. Yet, organ-specific mechanisms linking lymphatic dysfunction to disease etiology remain largely unknown. In this study, we uncover a novel intestinal lymphatic program that is linked to the left-right (LR) asymmetric transcription factor *Pitx2*. We show that deletion of the asymmetric *Pitx2* enhancer, *ASE*, alters normal lacteal development through the lacteal-associated contractile smooth muscle lineage. *ASE* deletion leads to abnormal muscle morphogenesis induced by oxidative stress, resulting in impaired lacteal extension and defective lymphatic-dependent lipid transport. Surprisingly, activation of lymphatic-independent trafficking directs dietary lipids from the gut directly to the liver, causing diet-induced fatty liver disease. In summary, our studies reveal the molecular mechanism linking gut lymphatic development to the earliest symmetry-breaking *Pitx2* and highlight the important relationship between intestinal lymphangiogenesis and gut-liver axis.

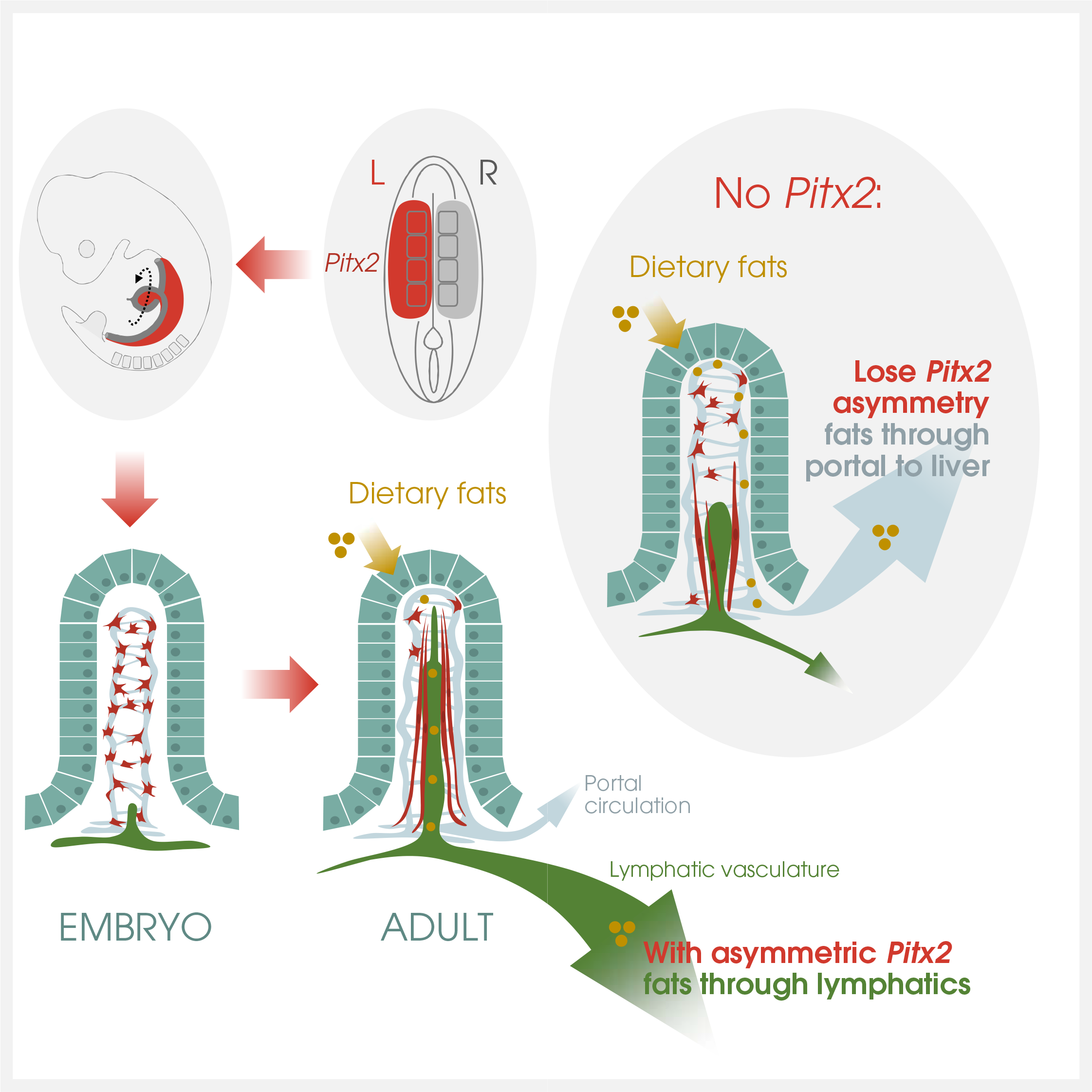
GRAPHICAL ABSTRACT

**HIGHLIGHTS:** ∼ Gut lymphangiogenesis is linked to *Pitx2*-driven LR asymmetry

∼ Lacteal-associated smooth muscle requires *ASE*

∼ *ASE* deletion leads to redox imbalance in intestinal smooth muscle lineage

∼ *ASE* is required for the normal route of dietary lipid transport

∼ *Pitx2^ASE/ASE^* neonates develop diet-induced fatty liver disease

## INTRODUCTION

Lymphatic vessels are a crucial component of the cardiovascular system and play a central role in several pathogenic processes including edema, hypertension, obesity, diabetes, inflammation, and metastasis [1–9]. In the intestine, the lymphatic vessels are essential for the absorption and transport of fats and fat-soluble nutrients, a function that distinguishes them from all other lymphatic channels. However, the molecular mechanisms governing their specialized functions remain unknown. Knowledge of how gut lymphatic development is regulated at the organ level will provide important insights into the mechanisms underlying these diseases.

The formation of gut lymphatic vessels follows blood vascular development [10]. Specifically, we have previously shown that gut lymphangiogenesis depends on prior establishment of arteries within the left side of the dorsal mesentery, the major conduit for blood and lymphatic vessels in the gut [11]. This process is regulated by the homeodomain transcription factor *Pitx2*, the major player responsible for the evolutionarily conserved left-right (LR) asymmetry of visceral organs [12–15]. Expression of *Pitx2* on the left side of the dorsal mesentery drives gut rotation and the resulting gut chirality. Disruption of this leads to intestinal malrotation and volvulus, a catastrophic strangulation of the gut vasculature [16–18]. In this context, gut rotation must be carefully coordinated with the early positioning of blood and lymphatic vessels to prevent vascular occlusion and to endow lymphatics with their polarized absorptive function once they arise in the intestinal villus. Indeed, gut arterial development is commensurate with the onset of gut rotation and proceeds strictly on the left side of the dorsal mesentery dependent on *Cxcl12*-*Cxcr4* signaling, downstream of Pitx2 [11]. Subsequent gut lymphangiogenesis is also left-sided and depends on prior arteriogenesis [11]. *Pitx2^-/-^* mouse embryos have gut arterial patterning defects and fail to initiate gut lymphatic development [11]. Thus, proper gut lymphatic development, like asymmetric gut rotation morphogenesis, depends on *Pitx2* to link vessel patterning with the morphogenesis of the parent organ.

The Pitx gene family includes three vertebrate paralogues, Pitx1, Pitx2, and Pitx3, that share an almost identical homeodomain protein sequence, varying mainly in the N-terminal region [19, 20]. In mice, the *Pitx2* gene maps to chromosome 3 [19] and is transcribed into three distinct isoforms: *Pitx2a, Pitx2b, and Pitx2c*. The *Pitx2a* and *Pitx2b* splice variants share the same promoter and are expressed bilaterally in the embryo, while *Pitx2c*, the major gut isoform, is transcribed from a separate promoter and is expressed asymmetrically [21–24]. Importantly, left-specific expression of *Pitx2c* is conferred by the evolutionarily conserved asymmetric *Pitx2* enhancer, *ASE* [21, 22]. *Pitx2^ASE/ASE^* mutant mice fail to manifest most of the left-sided *Pitx2c* expression and exhibit some laterality defects similar to those of *Pitx2^-/-^* mice [22, 25–27] and mice specifically lacking *Pitx2c* [28]. Importantly, unlike *Pitx2^-/-^* embryos, most *ASE* mutants survive to birth and some reach weaning age, allowing analyses of gut lymphatic function in the absence of *ASE*.

To analyze the consequence of *ASE* deletion, we first focused on the morphogenesis of lacteals, the absorptive lymphatic capillaries responsible for dietary fat uptake and gut immune surveillance [5, 29]. Fatty nutrients use lacteals as their major transport route, where lipids packaged into lipoprotein particles (chylomicrons) drain into larger mesenteric collecting vessels and ultimately to the systemic blood circulation [9, 30–32].

In contrast, non-lipid nutrients enter the portal circulation for primary delivery to the liver [9, 30–32]. Lacteals form around embryonic (E) day 17.5 and are functionally ready to absorb lipids from milk at birth [10]. Signaling via the lymphangiogenic vascular endothelial growth factor C (VEGF-C), the main VEGF receptor, VEGFR-3, and its co-receptor neuropilin 2 (NRP2), plays a key role in driving postnatal lacteal sprouting and lacteal growth [33, 34]. Although the majority of adult lymphatic vessels are quiescent, intestinal lacteals, similar to intestinal epithelium, are in a constant state of regeneration throughout adulthood, a process dependent on the formation of lacteal filopodia extensions at the lacteal tip [35]. Moreover, whereas most lymphatic capillaries lack smooth muscle coverage, lacteals are surrounded by contractile bundles of axial smooth muscle cells (SMCs) [36], which secrete lymphangiogenic growth factors [37] and facilitate lipid transport by squeezing the lacteal under autonomic nervous system control [38].

Here we show that *Pitx2* is required for the development and function of lacteals through a non-cell autonomous pathway that involves lacteal-associated contractile axial SMCs. Loss of asymmetric *Pitx2* expression, induced by *ASE* deletion, results in abnormal axial SMC morphogenesis caused by oxidative stress, leading to impaired extension of the developing lacteals and defective lymphatic-dependent lipid transport. Unexpectedly, we found that *ASE* deletion resulted in shunting of dietary lipid transport to the portal circulation resulting in excessive accumulation of lipids within the liver, and severe hepatic lipidosis (fatty liver disease) in *Pitx2^ASE/ASE^* neonatal pups. We propose that this alternative lymphatic-independent transport of dietary lipids is active in *Pitx2^ASE/ASE^* pups because lymphatic structure and transport within the intestinal villus is compromised. In summary, our studies reveal a molecular mechanism linking gut lymphatic development and function to the earliest symmetry-breaking events governed by *Pitx2* and highlight the important relationship between the intestinal lymphatic vasculature and the gut-liver axis. Moreover, our studies demonstrate how the *Pitx2* gene continues to orchestrate postnatal intestinal development and function beyond its well-established roles in early LR asymmetry.

## RESULTS

### Loss of LR asymmetric Pitx2 enhancer ASE results in growth retardation and early postnatal lethality

The *Pitx2* gene encodes three isoforms (*Pitx2a*, *Pitx2b, Pitx2c)* (Fig. 1A)*. Pitx2c*, the major isoform in the gut, is expressed asymmetrically via a conserved enhancer, *ASE* [22]. *Pitx2^-/-^* mouse embryos lacking all isoforms (*Pitx2*^hd/hd^) [25] fail to initiate gut lymphangiogenesis and die by E14.5 precluding analysis of gut lymphatic development [11] (Fig. 1BC). To explore the role of Pitx2 in gut lymphatics, we studied mice lacking *ASE* (*Pitx2^ASE/ASE^)*, which fail to manifest left-sided *Pitx2c* expression but survive to birth [22] (Fig. 1BC).

**Figure 1.**
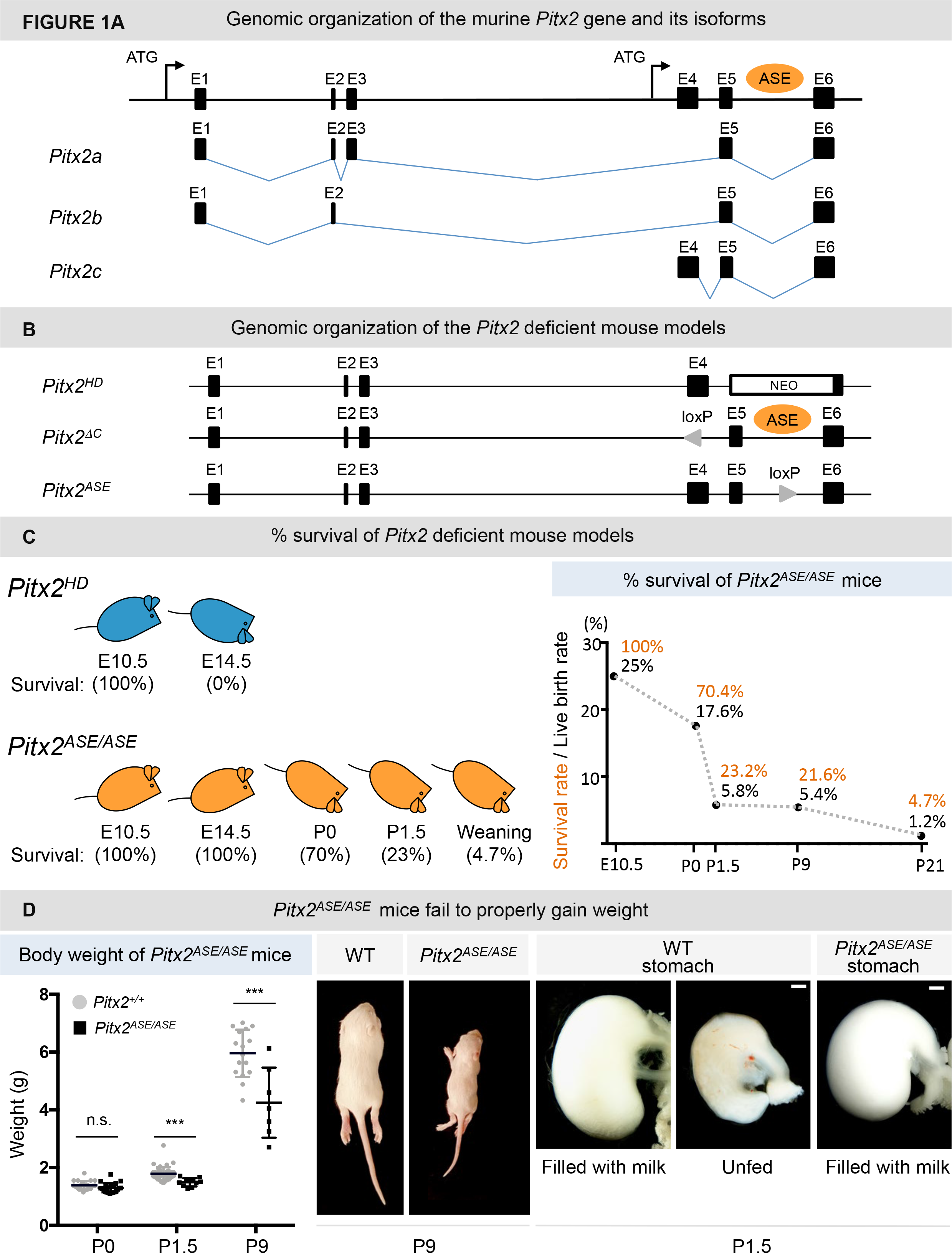
*Pitx2^ASE/ASE^* mice fail to properly gain weight. (A) The *Pitx2* gene structure and differential splicing of three *Pitx2* isoforms. E: exon; ATG: start codon; black arrow: transcriptional start site; ASE: <u>as</u>ymmetric <u>e</u>nhancer. (B) The genetic organization of *Pitx2* deficient mouse models. *Pitx2^HD^* mouse has <u>h</u>omeobox <u>d</u>eletion from exon 5 to 6. *Pitx2*^Δ^*^c^* (*Pitx2c*) mouse has *Pitx2c*-specific exon 4 deletion. *Pitx2^ASE^* mouse contains *ASE* deletion in intron 5. (C) Survival rates of *Pitx2^ASE/ASE^* mice at different stages. Data displayed on Y-axis are represented as incidence of live *Pitx2^ASE/ASE^* mice upon collection. Data calculated from the following pool of mice: 28 at E10.5, 125 at P0, 500 at P1.5, 533 at P9, and 169 at P21 (weaning age). Orange characters mark the survival rate of *Pitx2^ASE/ASE^* mice; calculations are based on the number of *Pitx2^ASE/ASE^* live mice divided by the expected number of mice according to Mendelian ratio (25%). Black characters mark live birth rate of *Pitx2^ASE/ASE^* mice; calculations are based on the number of *Pitx2^ASE/ASE^* live mice divided by number of total live mice. (D) *Pitx2^ASE/ASE^* mice gain weight less efficiently than their WT and heterozygous littermates. <u>Left</u>: Body weights measured at E18.5 and/or at birth are counted as P0. Body weights measured at P8.5 are counted as P9. <u>Middle</u>: A pair of littermates showing significant growth retardation of *Pitx2^ASE/ASE^ versus* WT mice at P9. <u>Right</u>: Stomach at collection of P1.5 WT fed, WT unfed, and *Pitx2^ASE/ASE^* fed littermates showing comparable food ingestion of mutant and WT mice. Data displayed on Y-axis in the left side graph are represented as mean ± SEM. P0 WT = 1.388 ± 0.03382 g, n = 21; P0 *Pitx2^ASE/ASE^* = 1.295 ± 0.03531 g, n = 22. Difference between WT and *Pitx2^ASE/ASE^* = 0.09264 ± 0.04897 g, P = 0.0656. P1.5 WT = 1.791 ± 0.2251 g, n = 41; P1.5 *Pitx2^ASE/ASE^* = 1.495 ± 0.1435g, n = 10. Difference between WT and *Pitx2^ASE/ASE^* = -0.2960 ± 0.07494g, P = 0.0003***. P9 WT = 5.963 ± 0.8161 g, n = 16; P9 *Pitx2^ASE/ASE^* = 4.25 ± 1.211 g, n = 7. Difference between WT and *Pitx2^ASE/ASE^* = -1.713 ± 0.4286 g, P = 0.0007***. Scale bar, 500 μm. See also Figure S1.

Whereas most *Pitx2^ASE/ASE^* mice die within hours after birth (C57BL/6 x 129 mixed background), we noted that some *Pitx2^ASE/ASE^* newborns on the FVB background remain viable for ∼10 days after birth (21.6%, Fig. 1C and Fig. S1) while an even smaller subset survive past weaning age (4.7%, Fig. 1C). Collectively, the *Pitx2^ASE/ASE^* mouse model allows analyses of gut lymphatic development and function in the absence of *ASE*.

Intestinal lymphatics are the essential channels for absorption and transport of dietary lipids. Thus, to briefly evaluate intestinal lymphatic function of *ASE* in dietary lipid transport, we first compared post-feeding weight gain between wild type (WT), *Pitx2^ASE/+^* heterozygous, and *Pitx2^ASE/ASE^* homozygous neonates. Despite being born with comparable body weight, *Pitx2^ASE/ASE^* homozygous pups had severely retarded growth at 24-48 hours and at postnatal (P) day P9 (Fig. 1D and Fig. S1). Importantly, this was not a consequence of variable food ingestion, as ample milk was seen in all neonatal stomachs analyzed, regardless of genotype (Fig. 1D). Thus, failure to properly gain weight in *Pitx2^ASE/ASE^* mutant pups was due to malabsorption rather than insufficient food intake, implicating a role for *ASE* (and *Pitx2c*) in lymphatic transport function.

### ASE is required for lacteal extension and formation of lacteal filopodia

Intestinal lipid transport is driven by lacteals at the center of each intestinal villus (Fig. S2A). We hypothesized that failure to properly gain weight in *Pitx2^ASE/ASE^* pups is due to abnormal lacteal development in the absence of *ASE*. Lacteals can be detected as early as E17.5 using the lymphatic endothelial cell marker Lyve-1 [10] (Fig. S2A). We first characterized embryonic and postnatal lacteal development in WT and *Pitx2^ASE/ASE^* mice using Lyve-1 immunofluorescence (IF) on gut slices (Fig. S2B). First, we observed a significant reduction in the number of Lyve-1+ lacteals in *Pitx2^ASE/ASE^* embryos at E18.5 (Fig. S2C) and again at P9 (Fig. S2C), suggesting a role for *Pitx2* in lacteal development. Next, we noted that lymphatic endothelial cells at the tip of the lacteals extend long filopodia in WT neonatal intestines, consistent with active sprouting and migration (P1.5, Fig. 2A, arrows) [39]. Moreover, extension of long filopodia was largely restricted to the tip cell of the lacteals (Fig. 2A, arrows). We observed that ∼ 73% of WT lacteals harbor at least one filopodium at P1.5 (Fig. 2A; 72.90 ± 2.760%, n=16), in agreement with prior studies in adulthood [35]. By contrast, we observed that most *Pitx2^ASE/ASE^* mutant lacteals are missing filopodia at P1.5 (Fig. 2A, arrowheads; 41.28 ± 3.545%, n=10, p=0.0001). Similar results were obtained following isoform-specific deletion of *Pitx2c* (Fig. 2B; WT: 81.38 ± 4.660, n=3; *Pitx2c^-/-^*, 41.13 ± 5.604, n=4; p=0.0034). Filopodia formation remained suppressed during later stages of development (P9), a phenotype that exacerbated with time (Fig. 2C; WT, 49.66 ± 4.200%, n=8; *Pitx2^ASE/ASE^*, 8.543 ± 2.610%, n=3; p=0.0003).

**Figure 2.**
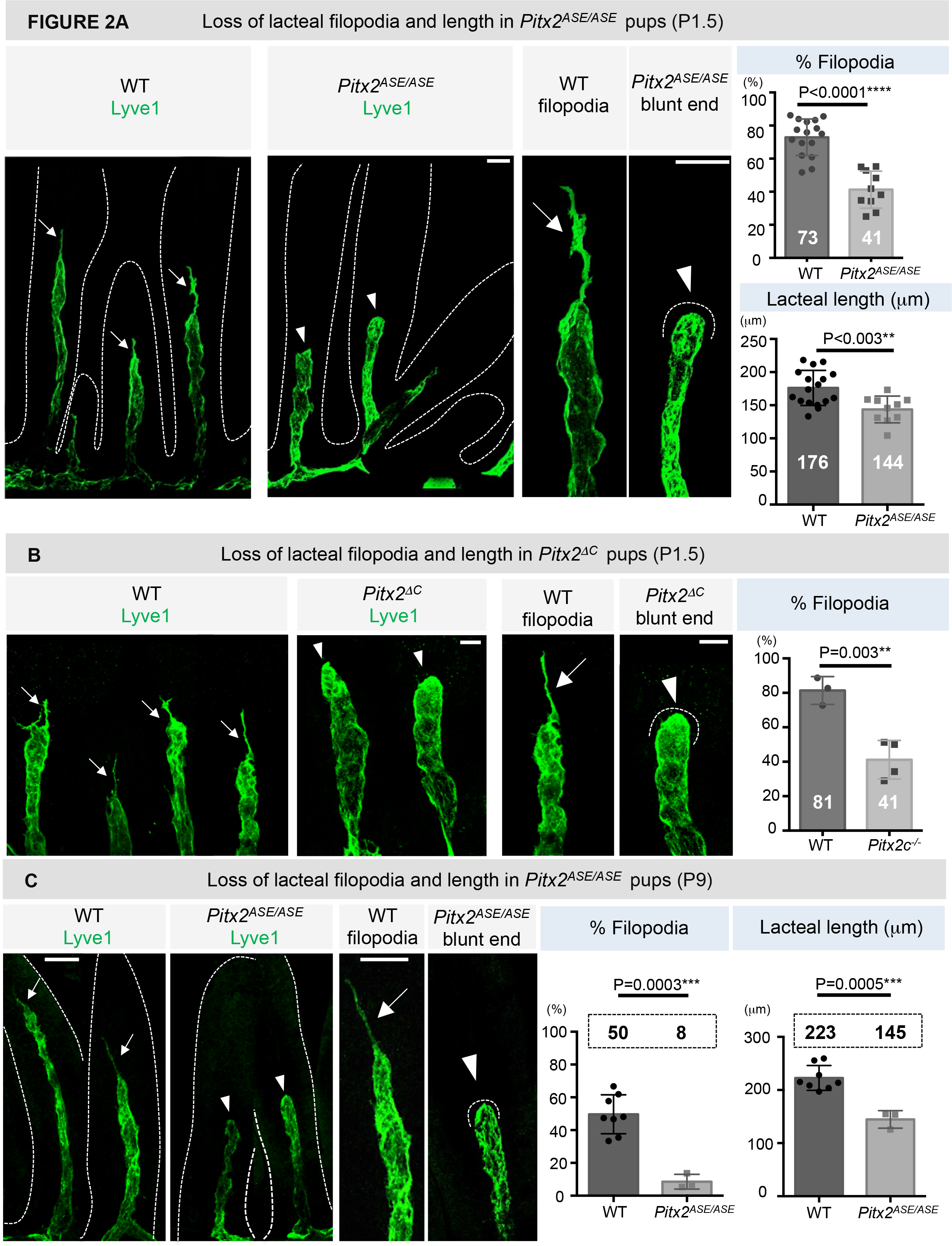
Shortened lacteals with missing filopodia in *Pitx2^ASE/ASE^* mice. (A) Representative images of whole mount lacteals (lyve-1, green) of WT and *Pitx2^ASE/ASE^* villi at P1.5. <u>Top right</u>: % lacteals with filopodia. <u>Bottom right</u>: Average lacteal length. Data are represented as mean ± SEM. P1.5 WT lacteals with filopodia = 72.90 ± 2.760 %, n = 16; P1.5 *Pitx2^ASE/ASE^* lacteals with filopodia = 41.28 ± 3.545 %, n = 10. Difference between WT and *Pitx2^ASE/ASE^* = -31.62 ± 4.476 %. P < 0.0001****. P1.5 WT lacteal length = 176.1 ± 6.401 μm, n=17; P1.5 *Pitx2^ASE^* lacteal length = 143.7 ± 6.392 μm, n = 10. Difference between WT and *Pitx2^ASE/ASE^* = -32.41 ± 9.704 μm. P = 0.0026**. White characters mark the mean in each group. Scale bars, 30 μm. (B) Representative images of whole mount lacteals (lyve-1, green) of WT and *Pitx2^ΔC^* villi at P1.5. <u>Right</u>: % lacteals with filopodia. Data are represented as mean ± SEM. WT = 81.38 ± 4.660 %, n = 3; *Pitx2^ΔC^* = 41.13 ± 5.604 %, n = 4; Difference between WT and *Pitx2^ΔC^* = -40.25 ± 7.692 %, P = 0.0034**. White characters mark the mean in each group. Scale bar, 20 μm. Representative images of whole mount lacteals (lyve-1, green) of WT and *Pitx2^ASE/ASE^* villi at P9. <u>Left</u>: % lacteals with filopodia. <u>Right</u>: Average lacteal length. Note the extended filopodium in WT lacteal tips (white arrows) vs. the blunt-ended lacteals (white arrowhead) in mutants. Data are represented as mean ± SEM. P9 WT lacteals with filopodia = 49.66 ± 4.200 %, n = 8; P9 *Pitx2^ASE/ASE^* lacteals with filopodia = 8.543 ± 2.610 %, n = 3. Difference between WT and *Pitx2^ASE/ASE^* = -41.12 ± 7.237 %. P = 0.0003***. P9 WT lacteal length = 222.7 ± 8.252 μm, n = 8; P9 *Pitx2^ASE/ASE^* lacteal length = 144.6 ± 9.518 μm, n = 3. Difference between WT and *Pitx2^ASE/ASE^* = -78.12 ± 14.90 μm. P = 0.0005***. Black characters mark the mean in each group. Scale bars, 30 μm. See also Figure S2.

Consistent with filopodia defects, *Pitx2^ASE/ASE^* lacteals were significantly shorter than those of control littermates at P1.5 (Fig. 2A; WT, 176.1 ± 6.401μm, n=17; *Pitx2^ASE/ASE^*, 143.7 ± 6.392μm, n=10, p= 0.0026) and again at P9 (Fig. 2C; WT, 222.7 ± 8.252μm, n=8; *Pitx2^ASE/ASE^*, 144.6 ± 9.518μm, n=3, p=0.0005). Whereas WT lacteals elongated by ∼ 50μm from P1.5 to P9 (Fig. 2AC; P1.5, 176.1 ± 6.401μm, n=17; P9, 222.7 ± 8.252μm, n=8), such lacteal extension was halted in *Pitx2^ASE/ASE^* mutant pups, despite normal villus length (Fig. 2AC; P1.5, 143.7 ± 6.392μm, n=10; *P9,* 144.6 ± 9.518μm, n=3). In summary, we conclude that during gut lymphatic development, *ASE* is required for lacteal extension and lacteal filopodia formation.

### Pitx2 daughter cells populate lacteal-associated smooth muscle

Whereas a vascular-specific role for *Pitx2* has been documented both *in vitro* and *in vivo* [11, 22, 40–42], to our knowledge, endothelial cell-specific expression of *Pitx2* has never been detected. To determine whether *Pitx2* acts autonomously in lymphatic endothelial cells or in a non-cell autonomous manner during lacteal morphogenesis, we performed *Pitx2^Cre^* (knock-in) lineage tracing to follow *Pitx2* daughter cells in the gut (Fig. S3AA’A”). *Pitx2^Cre^* is a null *Pitx2* allele that drives Cre activity in cells that have expressed any *Pitx2* isoform throughout embryonic development, including the *Pitx2c* asymmetric isoform [28] (Fig. S3A). To follow *Pitx2* daughter cells, we crossed *Pitx2^cre^* heterozygotes with *Rosa26* (*R26R-tdTomato*) reporter mice (Fig. S3A).

TdTomato-marked *Pitx2* daughter cells were easily detected in the intestine, consistent with the described role of *Pitx2* during midgut development (Fig. S3A’). Consistent with prior reports, *Pitx2* daughter cells did not populate CD31-positive blood endothelial cells (BECs), or Lyve1-positive lymphatic endothelial cells (LECs) at any time point analyzed (E14.5, E16.5, E18.5, P1.5, P9, 6 months, n=19) (Fig. S3A”). Instead, we found *Pitx2* daughter cells among the α-smooth muscle actin (αSMA)-positive smooth muscle cells of the intestine, including villus-axial smooth muscle cells (SMCs) that are closely associated with lacteals [36, 43] (Fig. 3A’A”A”’ white arrowheads and Fig. 3B, WT). We confirmed this observation using *ASE*-specific lineage tracing with *Rosa26* (*R26R-EYFP*) reporter mice, where the activity of Cre is driven by an ∼18kb genomic fragment containing *ASE* and the *Pitx2c* promoter [44] (Fig. 3A””, white arrowheads).

**Figure 3.**
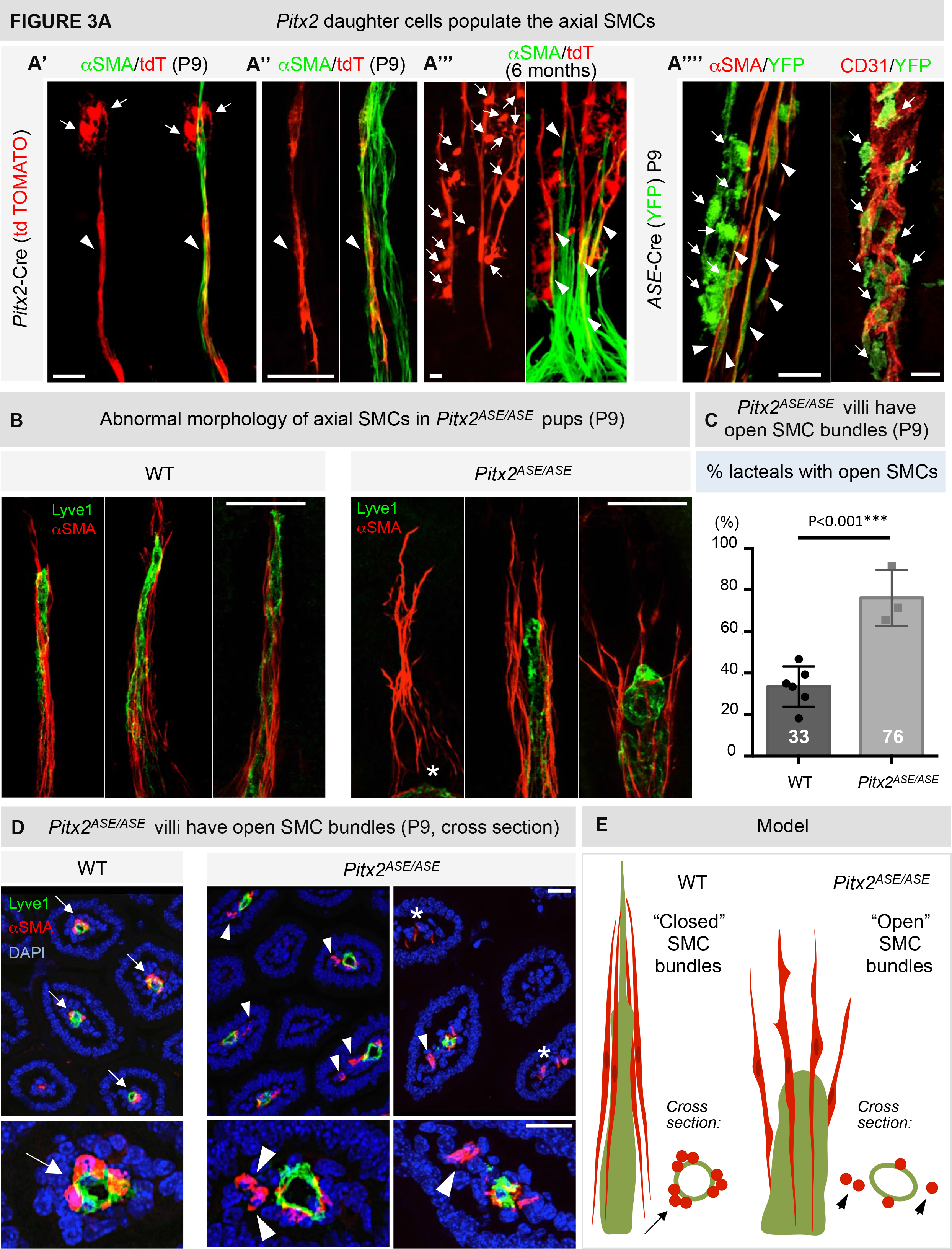
Malformation of the muscular-lacteal complex in *Pitx2^ASE/ASE^* mice. (A) Lineage tracing using *Pitx2^cre^* - *ROSA26^CAG-tdTomato^* and *ASE^cre^* - *ROSA26^CAG-EYFP^* mice. Representative image of whole mount villus from Pitx2*^cre^*:: *ROSA26^CAG-tdTomato^* mice at P9 (A’, A”) and 6 months (A”’). Note the colocalization of αSMA (green) and tdTomato (tdT, red) that marks axial SMCs of *Pitx2* descend (white arrowhead) and axial SMC heterogeneity (A”). White arrows depict additional cell types marked by *Pitx2* daughter cells (A’). (A””) Representative images of whole mount villus from *ASE^cre^* - *ROSA26^CAG-EYFP^* mice at P9. Note the colocalization of αSMA (red) and YFP (green) that marks axial SMCs of *ASE* descent (white arrowheads). White arrows depict additional cell types marked by *ASE* daughter cells, distinct from BECs (red). Scale bar, 20 μm. (B) Representative images of whole mount muscular-lacteal complexes of WT and *Pitx2^ASE/ASE^* mice at P9. Green is lacteal (lyve-1); red is αSMA (smooth muscle). From left to right in the *Pitx2^ASE/ASE^* panel: aberrant muscular arrangement with no lacteal (white asterisk); loss of direct contact between axial SMCs from the lacteal; opened axial SMCs with a shorter and dilated lacteal. Scale bar, 40 μm. (C) % lacteals surrounded by “open” axial SMCs based on data represented in (C). An axial muscle structure was characterized by the apical part arrangement. We defined the muscles “opened” if the axial muscles were divergently aligned around the lacteal tip as demonstrated in (F). Data are represented as mean ± SEM. P9 WT = 33.51 ± 3.950 %, n = 6; P9 *Pitx2^ASE/ASE^* = 76.12 ± 7.775 %, n = 3. Difference between WT and *Pitx2^ASE/ASE^* = 42.61 ± 7.703 %, P = 0.0009***. White characters mark the mean in each group. (D) Representative images of muscular-lacteal complexes of WT and *Pitx2^ASE/ASE^* mice at P9 in transverse sections. Green is lacteal (lyve-1); red is αSMA (smooth muscle); blue is DAPI. Note the close (white arrows) vs. distanced (white arrowheads) proximity of red (muscle) and green (lacteal) signals. White stars mark villi with missing lacteals. Scale bar, 20 μm. (E) Cartoon model demonstrating muscular-lacteal arrangement of WT and *Pitx2^ASE/ASE^* mice. Green represents lacteal; red is smooth muscle. See also Figure S3.

In addition to axial SMCs, we found a discrete fraction of TdTomato-marked *Pitx2* (Fig. 3A’, white arrows) and YFP-marked *ASE* (Fig. 3A””, white arrows) daughter cells morphologically different from axial SMCs and in close proximity to, but distinct from, the CD31+ blood capillary plexus of the villous lamina propria (Fig. 3A””, white arrows). Based on their location within the villus and morphological appearance these cells may lie upstream in the lineage compared to the more mature axial SMCs and may be involved in muscular regeneration at later stages (Fig. 3A”’, white arrows, 6 months). Taken together, these findings suggest that *Pitx2* regulates lacteal development through a non-cell autonomous pathway, and underscore a role for *Pitx2* in smooth muscle patterning of the intestinal lamina propria.

### Loss of ASE impairs lacteal-associated axial smooth muscle development

A subset of axial SMCs secretes VEGF-C [37], the critical lymphatic morphogen. Loss of VEGF-C-driven activation of VEGFR-3, the main VEGF receptor expressed by lacteals, leads to arrested lacteal growth [37]. Moreover, axial SMC contraction can squeeze the adjacent lacteal to drive lipid transport [38]. Therefore, villus-axial SMCs may provide the principal cues for development of lacteals. We therefore hypothesized that *Pitx2* is indirectly required for lacteal morphogenesis via SMC patterning. We first studied the villus smooth muscle histopathological structure in WT and *Pitx2^ASE/ASE^* mice post weaning, at P26 (Fig. S3B). Compared to robust villus smooth muscle coverage in WT mice, we observed a severe reduction in smooth muscle of the lamina propria in *Pitx2^ASE/ASE^* mice, consistent with a role of *Pitx2* in villus smooth muscle development (Fig. S3B, red arrowheads).

To corroborate these findings and to characterize the muscular-lacteal complex inside the villus at high resolution, we examined WT and *Pitx2^ASE/ASE^* gut slices using confocal microscopy with αSMA (smooth muscle) and Lyve-1 (lacteal) IF. In WT pups at P9, αSMA+ cells were closely intertwined with each other and with Lyve-1+ lacteals, a striking pattern with “closed” muscle bundle morphology, suggesting active muscular contraction squeezing the lacteal (Fig. 3BE). In contrast, age-matched *Pitx2^ASE/ASE^* mutants displayed a wide spectrum of muscular-lacteal abnormalities, including misaligned axial SMCs with absent lacteals, reduced axial SMC numbers, loss of direct SMCs-LECs contact, and “open” bundle conformation leaving spaces between individual muscle bundles and lacteals (Fig. 3BE). We quantified a significant increase in open axial smooth muscle bundles within *Pitx2^ASE/ASE^* villi, compared with control mice (Fig. 3BC; WT, 33.51 ± 3.950%, n=6; *Pitx2^ASE/ASE^*, 76.12 ± 7.775%, n=3, p=0.0009). Moreover, using immunohistochemistry (IHC) of transverse tissue sections through the villus, we showed that unlike the close association of WT axial SMCs with adjacent lacteals (Fig. 3DE, arrows), *Pitx2^ASE/ASE^* axial SMCs and lacteals failed to tightly align (Fig. 3DE, arrowheads). Thus, *ASE* regulates the morphogenesis of villus axial smooth muscle, which further directs intestinal lymphatic development.

### Lacteal-associated axial SMCs arise by ASE-dependent remodeling

Villus-axial SMCs were first identified in ultrastructural studies in 1909 [45]. The ultrastructure was characterized in the rat and was considered vital to providing structural tension against interstitial pressure while driving villous rhythmical contraction in close relationship with the adjacent fibroblasts [36, 46]. More recent *in vivo* imaging in mice has revealed that villus SMCs promote lateral villus contractions, dependent on the autonomic nervous system, and facilitate lipid transport by squeezing the lacteal [38]. Despite the crucial interplay between axial SMCs and lacteal function, the origins and patterning mechanisms of axial SMCs remain unexplored.

To investigate muscle formation alongside lacteals, we assessed the dynamics of villous muscular formation in WT mice from E16.5 to P9 (Fig. 4A and Fig. S4A). Cells expressing low levels of αSMA (αSMA+) were first detected in the villus at E16.5, before the emergence of lacteals (Fig. S4A). Morphologically, αSMA+ cells within the villus appeared star-shaped with multiple protrusions (Fig. 4A, inset, “star cells”), and aligned closely with the CD31+ blood vascular plexus (Fig. S4A). Interestingly, αSMA+ star cells were reminiscent of TdTomato-marked *Pitx2* and YFP-marked *ASE* daughter cells we previously observed (Fig. 3A’A””, white arrows). Strikingly, coincident with the appearance of lacteals at E18.5, αSMA+ cells underwent a robust transformation to elongated, spindle-like cell types with strong immunoreactivity for αSMA (αSMA++), and in close association with lacteals (Fig. 4A, graph). At P1.5, the number of spindle-like αSMA++ cells became predominant at the expense of αSMA+ star cells (Fig. 4A, graph). At P9, all αSMA++ spindle cells were located adjacent to the lacteal, extending radially from the submucosa to the filopodium at the tip of the lacteal, while very few αSMA+ star cells were detected at this time (Fig. 4A and Fig. S4D).

**Figure 4.**
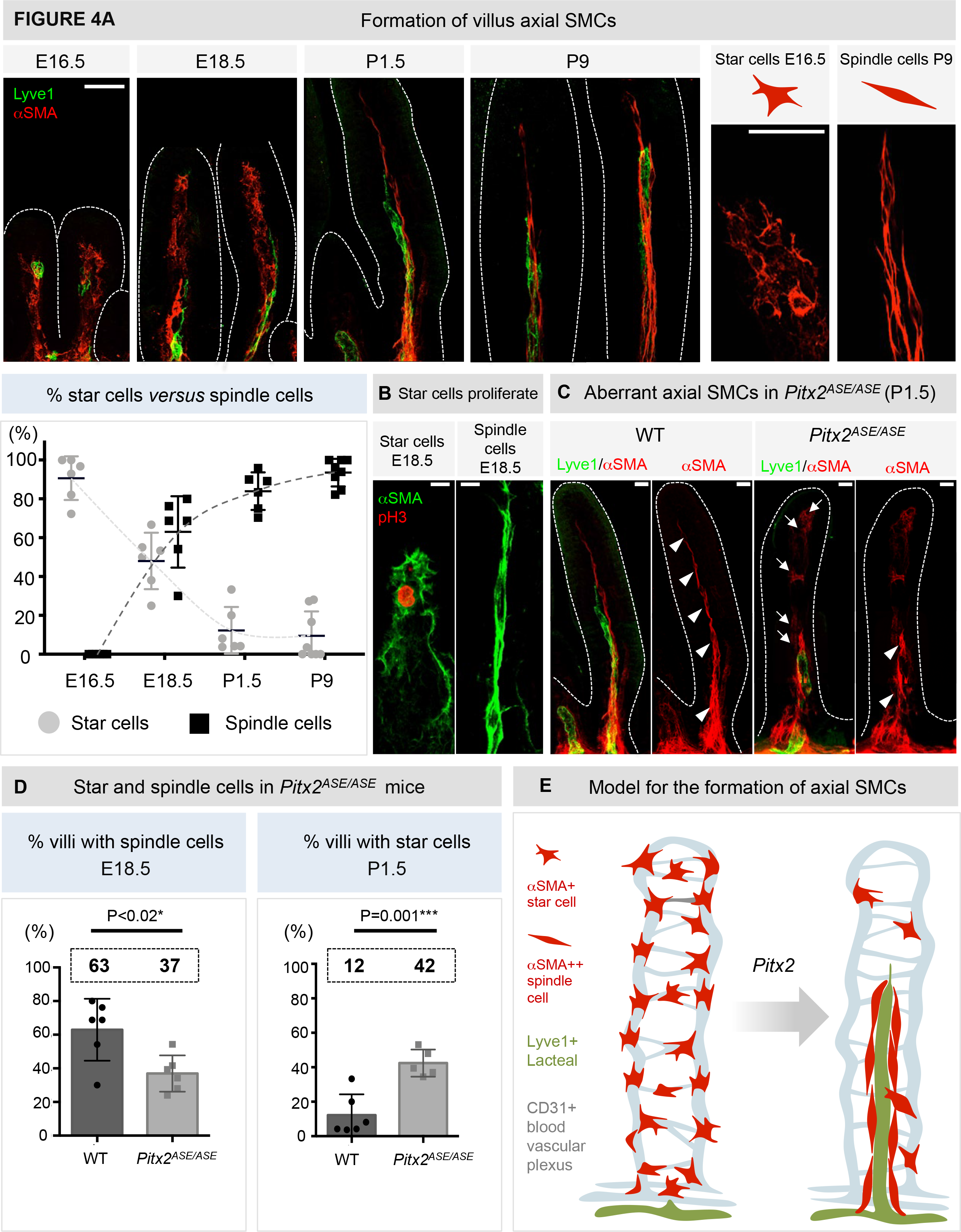
Lacteal-associated axial SMCs arise by *ASE*-dependent remodeling. (A) Representative images of whole mount WT villous muscular-lacteal development from E16.5-P9. Green is lacteal (lyve-1), red is αSMA (smooth muscle). Note the interaction of muscle (red) with the developing lacteal (green). <u>Insets</u>: Note the shape difference of cells at E16.5 (star cells) *versus* at P9 (spindle cell). Scale bar, 40 μm. <u>Graph:</u> % villi with αSMA+ star cells (black squares) or αSMA++ spindle cells (gray dots) during development. Note the negative correlation between αSMA+ star cells (dominant in prenatal stages) and αSMA++ spindle cells (dominant postnatally). Data displayed on Y-axis are represented as mean ± SEM. E16.5 villi with star cells = 90.7 ± 4.6 %, n = 6; E16.5 villi with bundles = 0 ± 0 %, n = 6. E18.5 villi with star cells = 48.02 ± 5.9 %, n = 6; E18.5 villi with muscle bundles = 62.98 ± 7.5 %, n = 6. P1.5 villi with star cells = 12.25 ± 4.9 %, n = 6; P1.5 villi with muscle bundles = 83.95 ± 4.0 %, n = 6. P9 villi with star cells = 9.454 ± 4.5%, n = 8; P9 villi with muscle bundles = 93.51 ± 2.5 %, n = 8. (B) Star cells, but not spindle cells, are proliferative (E18.5). Representative images of whole mount WT villus; green is αSMA (green), red is phospho-histone 3 (H3P), and blue is DAPI. Note that pH3 signal colocalized only with star-like but not spindle-like SMCs. Scale bar, 10 μm. (C) Representative images of whole mount muscular-lacteal complexes from WT and *Pitx2^ASE/ASE^* siblings at P9. Green is lacteal (lyve-1), red is αSMA (smooth muscle). Note the retention of SMA^+^ star cells in *Pitx2^ASE/ASE^*. Scale bar, 20 μm. (D) Graphs demonstrating % villi with star or spindle cells in WT and *Pitx2^ASE/ASE^* siblings. <u>Left</u>: % villi with spindle cells in E18.5 embryos. <u>Right</u>: % villi with star cells in P1.5 pups. Data displayed on Y-axes are represented as mean ± SEM. E18.5 WT villi with spindles = 62.98 ± 7.52 %, n = 6; E18.5 *Pitx2^ASE/ASE^* villi with spindles = 36.93 ± 4.41 %, n = 6. Difference between WT and *Pitx2^ASE/ASE^* = -26.05 ± 8.72 %, P = 0.0136*. P1.5 WT villi with stars = 12.25 ± 4.917 %, n = 6; P1.5 *Pitx2^ASE^* villi with stars = 42.42 ± 3.510 %, n = 5. Difference between WT and *Pitx2^ASE/ASE^* = 30.18 ± 6.292 %, P = 0.0010***. (E) Cartoon model depicting star-to-spindle smooth muscle transition underlines the formation of axial SMC-lacteal complex. See also Figure S4.

Interestingly, αSMA+ star cells were actively proliferative (Fig. 4B, ∼ 10%) compared to very few proliferating αSMA++ spindle-like axial SMCs (Fig. 4B), as judged by phospho-histone H3 (pH3) immunoreactivity. This agrees with previous observations that only round muscular progenitor cells (but not mature spindle muscles) are proliferative [47, 48]. Consistent with αSMA++ spindle cells resembling more differentiated muscle cells, they expressed high levels of myosin heavy chain 11 (Myh11), one of the major contractile proteins in SMCs [49] (P9, Fig. S4D). In contrast, (putative progenitor) αSMA+ star cells showed very weak immunoreactivity for Myh11 (P9, Fig. S4D, white arrows). Collectively, these observations suggest that villus-axial smooth muscle development is commensurate with the formation of lacteals and proceeds by a robust spatiotemporal remodeling program whereby αSMA+ precursor star cells proliferate and assemble to create and shape the mature axial SMC network of the lacteal (Fig. 4E). Of note, axial SMCs were largely juxtavascular and this was especially evident for SMCs at the lacteal filopodia (Fig. S4C), highlighting an important reciprocal interaction with the adjacent lymphatic endothelium.

Next, we examined villus-axial SMC dynamics in the absence of *ASE*. As in WT villi, we first detected αSMA+ star cells in the *Pitx2^ASE/ASE^* mutants at E16.5, with no differences observed in the proportion of villi containing αSMA+ star cells among all genotypes tested (data not shown). However, two days later, at E18.5 we found fewer αSMA++ spindle-like axial SMCs in *Pitx2^ASE/ASE^* mutants compared to WT littermates (Fig. 4D; WT, 62.98 ± 7.52%, n=6; *Pitx2^ASE/ASE^*, 36.93± 4.41%, n=6, p=0.0136). Moreover, in contrast to WT neonates, a significant number of αSMA+ star cells remained in the lamina propria of *Pitx2^ASE/ASE^* mutants at P1.5 (Fig. 4CD; WT, 12.25 ± 4.917%, n=6; *Pitx2^ASE/ASE^*, 42.42 ± 3.510%, n=5, p= 0.0010). These star cell-retained *Pitx2^ASE/ASE^* villi (Fig. 4C, white arrows) had less prominent muscular bundles surrounding the lacteals (Fig. 4C, white arrowheads).

TUNEL staining of WT and *Pitx2^ASE/ASE^* mutants at P9 revealed no difference in villous SMC cell death (Fig. S4E). Using IHC and histopathological analyses, apoptotic or necrotic cells were not seen in E18.5, P1.5, and P9 *Pitx2^ASE/ASE^* villi, while villous SMC proliferation was similarly unaffected in *Pitx2^ASE/ASE^* at E18.5 (data not shown). These observations suggest that neither cell death nor cell cycle arrest at the stages examined contributes to the *ASE*-dependent axial SMC phenotypes. Instead, we propose that *ASE* functions in villous-axial SMC morphogenesis during their fetal to neonatal transition (Fig. 4E).

### *ASE* deletion leads to redox imbalance in intestinal smooth muscle

To explore the cause of muscular-lacteal impairment in *Pitx2^ASE/ASE^* mutant intestines, whole intestine transcriptomics (RNA-seq) of both WT and *Pitx2^ASE/ASE^* neonatal tissue at P1.5 was performed. Notably, ∼ 20% of genes differentially expressed between WT and *Pitx2^ASE/ASE^* mutant intestines corresponded to reactive oxygen species (ROS) production, antioxidant scavenging genes, and redox-sensitive signaling pathways (Fig. 5A). Among potential targets were genes that protect the cell from oxidative stress, such as the glutamate-cysteine ligase (*Gclc*, also known as gamma-glutamylcysteine synthetase), the first rate-limiting enzyme of glutathione synthesis [50–52]. Interestingly, half of the ROS-related differentially expressed genes are directly bound by Pitx2 *in vivo* [51] including the noncoding region of *Gclc* (Fig. S5A). These data raised the possibility that *Pitx2c* may directly regulate oxidation-reduction (redox) balance in intestinal smooth muscle.

**Figure 5.**
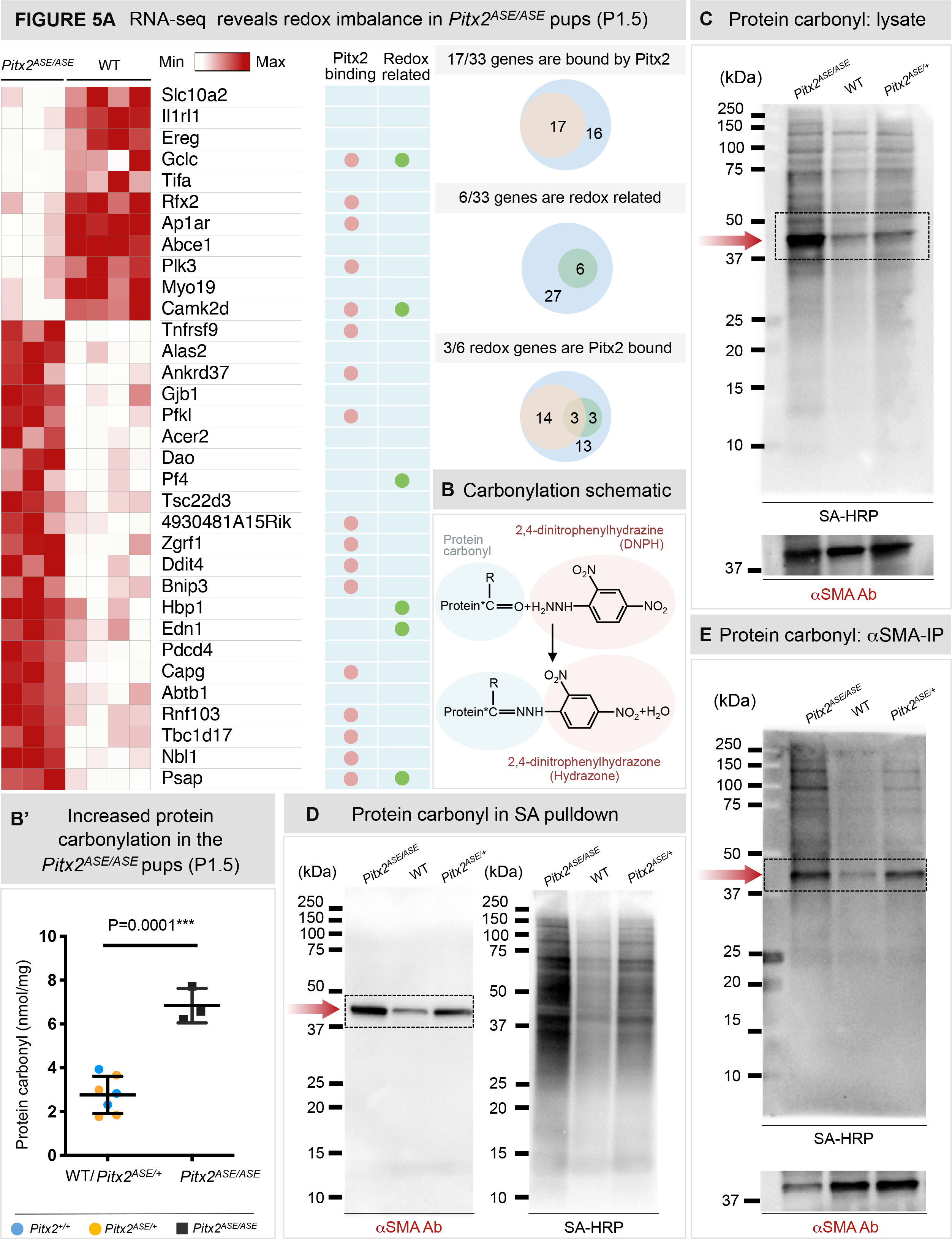
*ASE* deletion leads to redox imbalance and actin carbonylation in intestinal muscle. (A) RNA-seq data analysis. <u>Left</u>: Heatmap displaying the genes that were differentially expressed between WT and *Pitx2^ASE/ASE^* intestines at P1.5. The relative expression levels were displayed in color code based on the raw counts in each gene from each sample, with the minimum expression depicted in white and the maximum expression in dark red. Differentially expressed genes were identified as described in methods and listed in UCSC gene symbol. Genes bound by Pitx2 based on prior ChIP-seq were noted with red dots, genes with known function related to cellular redox homeostasis were noted with green dots. <u>Right</u>: Venn diagrams demonstrating the relation of differentially expressed genes bound by *Pitx2* and/or with reported function in cellular redox balance. (B) Schematic of protein carbonyl detection. 2,4-dinitrophenylhydrazine (DNPH, or hydrazine) reacts with protein carbonyl groups leading to the formation of the stable 2,4-dinitrophenylhydrazone (hydrazone) that can be detected and quantified spectrophotometrically (B’). Protein carbonyl concentration measured from WT (blue), *Pitx2^ASE/+^* (orange), and *Pitx2^ASE/ASE^* (black) intestines. Data are represented as mean ± SEM. Protein carbonyl in WT/ *Pitx2^ASE/+^* group = 2.76 ± 0.3194 nmol/mg, n = 7; protein carbonyl in *Pitx2^ASE/ASE^* = 6.833 ± 0.4567 nmol/mg, n = 3. Difference between WT/het and *Pitx2^ASE/ASE^* = 4.073 ± 0.5741 nmol/mg, P = 0.0001***. (C-E) Increased αSMA carbonyl level in *Pitx2^ASE/ASE^* intestines (P1.5). Representative blots of biotin-hydrazine (biotin-HZ) labeling assay demonstrate increased carbonylated αSMA in *Pitx2^ASE/ASE^* intestine. (C) P1.5 intestine protein lysates were labeled with biotin-HZ and separated by SDS-PAGE then transferred to nitrocellulose membrane. Total protein carbonyl was detected by Streptavidin-HRP (SA-HRP) (top blot), while total αSMA expression was detected by αSMA antibody on the same blot (lower blot). (D) P1.5 intestine protein extracts were incubated with biotin-HZ. Biotinylated proteins were then purified by streptavidin beads, separated by SDS-PAGE, and transferred to nitrocellulose membrane. (Left) αSMA was detected by αSMA antibody. (Right) Total protein carbonyl was detected by SA-HRP on the same blot. (E) P1.5 intestine protein extracts were labeled with biotin-HZ following αSMA immunoprecipitation (SMA-IP) and separated by SDS-PAGE then transferred to nitrocellulose membrane. Total protein carbonyl was detected by SA-HRP (top blot). SMA-IP efficiency was confirmed by total αSMA expression, detected by αSMA antibody on the same blot. kDa: kilo Dalton. See also Figure S5.

To investigate whether *ASE* deletion leads to a redox imbalance in the intestine, we detected and quantified spectrophotometrically protein carbonylation in WT, Pitx2*^ASE/+^* heterozygous, and Pitx2*^ASE/ASE^* neonatal intestines at P1.5. Carbonyl groups (Fig. 5B) are an irreversible product of ROS-mediated protein oxidation and a general biomarker for oxidative stress [52–54]. We observed Pitx2*^ASE/ASE^* intestines harbor 2.5-times more protein carbonyl content compared to their WT or Pitx2*^ASE/+^* littermates (Fig. 5B’; WT/Het: 2.760 ± 0.3194 nmol/mg, n=7; *Pitx2^ASE/ASE^* 6.833 ± 0.4567 nmol/mg, n=3, p=0.0001), demonstrating that *ASE* loss increases irreversible oxidative damage in the neonatal intestine. We also assessed protein carbonylation levels by electrophoresis, after labeling carbonyl groups in protein extracts from WT, *Pitx2^ASE/+^*, and *Pitx2^ASE/ASE^* neonatal intestines at P1.5 with biotin-hydrazine (biotin-HZ). Proteins were SDS-PAGE-separated, and probed on nitrocellulose with streptavidin-conjugated horseradish peroxidase (SA-HRP). Consistent with the spectroscopic measurements (Fig. 5B’), a higher total level of carbonylated proteins (overall darker lane signal) was observed in the Pitx2*^ASE/ASE^* intestine lysate relative to controls (Fig. 5C).

Interestingly, a well-defined carbonylated protein band of ∼ 42kDa was enriched in *Pitx2^ASE/ASE^* intestinal extracts versus controls (Fig. 5C). This band is consistent in size with monomeric actin. Probing the same lysates for αSMA showed actin levels were similar among all genotypes (Fig. 5C), suggesting the possibility that levels of oxidized actin were enriched in *Pitx2^ASE/ASE^* intestines.

The dynamic actin cytoskeleton is susceptible to oxidation, which regulates cell behavior and contractility pathways [55]. Oxidized actin is linked to decreased actin monomer (G-actin) assembly and actin filament (F-actin) stability [56]. Oxidation of select cysteine and methionine residues in actin causes structural changes, monomer aggregation, decreased actin polymerization and stability, and defective cell contractility [57–60]. Actin carbonylation is associated with even higher oxidant levels than those associated with the modification of critical cysteine and methionine residues, and an accumulation of carbonylated actin has been proposed to indicate severe oxidative stress and functional impairment [61]. Interestingly, we noted discontinuous patterns of αSMA arrangement in *Pitx2^ASE/ASE^* axial SMCs reminiscent of the severed F-actin filaments seen under oxidative stress (Fig. S5B). Altogether these data led us to hypothesize that *ASE* deletion impaired villus-axial smooth muscle morphogenesis due to abnormal accumulation of high levels of ROS in the differentiating smooth muscle.

To test our hypothesis, we directly assessed the oxidation status of smooth muscle actin in WT and *Pitx2^ASE/ASE^* neonatal intestines (Fig. 5D-E). Streptavidin-enrichment of biotin-labeled carbonylated proteins confirmed that actin was carbonylated and that *Pitx2^ASE/ASE^* yielded the highest amount of carbonylated αSMA (Fig. 5D). Immunoisolation of αSMA from biotin-HZ-labeled lysates similarly confirmed an increased level of carbonylated αSMA in *Pitx2^ASE/ASE^* intestine by comparison to WT or *Pitx2^ASE/+^* heterozygous littermates (Fig. 5E). These results demonstrate that loss of *ASE* induced irreversible oxidative damage in αSMA+ cells of the neonatal intestine. Collectively, our data suggest that elevated levels of oxidative stress caused muscular-lacteal impairment in *Pitx2^ASE/ASE^* intestines, pointing to a crucial role for *Pitx2* in the control of ROS during early smooth muscle development.

### *ASE* is required for the normal route of dietary lipid transport

The intestinal lacteal-lymphatic vasculature network is the major route through which long-chain fatty acids are absorbed as triglycerides into the body [62]. These lipids are packaged into large specialized lipoproteins called chylomicrons, which are secreted by intestinal epithelial cells [63, 64]. In contrast, non-fatty nutrients and shorter chain fatty acids are able to directly enter the hepatic portal circulation for processing within the liver [62]. This lymphatic-independent portal transport is insignificant compared with the lymphatic route, except under certain pathological conditions where disrupted intestinal circulation leads to lipid accumulation in the liver [65–74].

To assess whether the intestinal lymphatic structural defects caused by *ASE* deletion impaired gut lymphatic function, we orally administered fluorescent BODIPY C16, a fluorescent long-chain fatty acid tracer, to WT and *Pitx2^ASE/ASE^* neonatal mice to follow lipid trafficking (Fig. 6A) [75]. As expected, long-chain BODIPY C16 entered the lumen of the intestine and mesenteric collecting vessels after oral administration (Fig. 6A), with very little staining of hepatic portal circulation in WT pups (Fig. 6B). Strikingly, long-chain BODIPY C16 accumulation was found in the hepatic portal vein and liver parenchyma of *Pitx2^ASE/ASE^* mutant pups (Fig. 6B; liver, liver section, hepatic portal vein), confirming abnormal lipid trafficking in the absence of *ASE*. Notably, BODIPY C16 remained detectable in the mesenteric collecting lymph vessels of *Pitx2^ASE/ASE^* pups implying a partial, rather than complete, shift in lipid transport in the absence of *ASE* (Fig. S6). Collectively, *ASE* deletion impairs normal long-chain fatty acid lymphatic trafficking resulting in direct lipid entry into the hepatic portal circulation (Fig. 6C).

**Figure 6.**
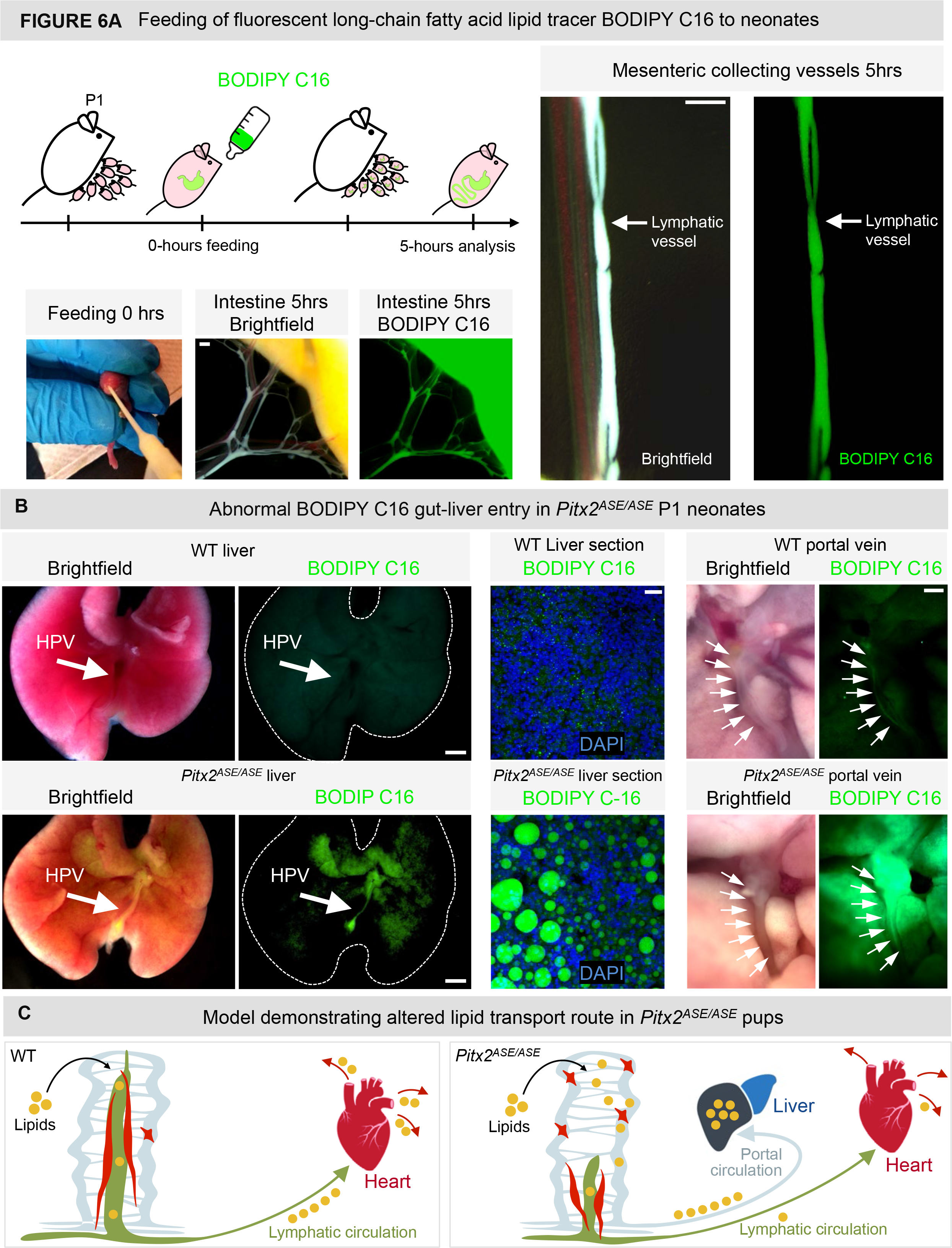
*ASE* is required for the normal route of dietary lipid transport. (A) Schematic demonstration of BODIPY™ FL C16 fluorescent lipid tracer feeding assay. Pups were fed with BODIPY™ FL C16 (long-chain fatty acid, green) then left with the dam for 5 hours before further analyses. Note representative images of mesenteric collecting lymphatic lipid tracer entry in WT neonates validating the feeding assay. Scale bar, 200 μm. (B) Hepatic analysis showing BODIPY™ FL C16 accumulation in *Pitx2^ASE/ASE^* at P1. <u>Left-to-right:</u> Analysis of whole livers, frozen liver sections with DAPI counterstain (blue), and hepatic portal vein (HPV, white arrows) in WT (top) and *Pitx2^ASE/ASE^* (bottom) neonates. Note enriched green signal in HPV and livers of *Pitx2^ASE/ASE^* neonates by comparison to their WT siblings. Scale bar, 1mm (left, liver whole organ), 20 μm (center, liver section), and 300 μm (right, HPV). (C) Cartoon model of lipid transport pathways in WT and *Pitx2^ASE/ASE^* neonates. Lymphatic-dependent transport (green) of dietary long-chain fatty acids (yellow dots) is dominant in WT mice, while additional activation of lymphatic-independent lipid transport via the hepatic portal vein (light blue) is evident in *Pitx2^ASE/ASE^* mice. Star cells and axial SMCs are shown in red. See also Figure S6.

### *Pitx2^ASE/ASE^* mice develop diet-induced fatty liver disease

Aberrant dietary lipid transport by the hepatic portal venous circulation can accumulate lipid within hepatocytes, resulting in histological features consistent with hepatic lipidosis. Indeed, changes characteristic of lipid accumulation were grossly apparent in *Pitx2^ASE/ASE^* mutant livers at P9. Thus, we compared liver lipid content in prenatal and postnatal WT and *Pitx2^ASE/ASE^* mice using oil red O, a fat-soluble dye that stains neutral triglycerides and lipids [76] (Fig. 7A). Consistent with our prior BODIPY C16 feeding data, increased oil red O staining in livers of postnatal, but not prenatal (unfed), Pitx2*^ASE/ASE^* mice, suggested hepatic lipid accumulation in *Pitx2^ASE/ASE^* livers is a consequence of post-partum nursing. Excessive hepatic lipid accumulation was observed as early as P1.5 and in 100% (n=4/4) of *Pitx2^ASE/ASE^* pups analyzed (Fig. 7A). Hepatic lipid accumulation in P1.5 was confirmed based on by ultrastructural examination by transmission electron microscopy (tEM) (Fig. S7A).

**Figure 7.**
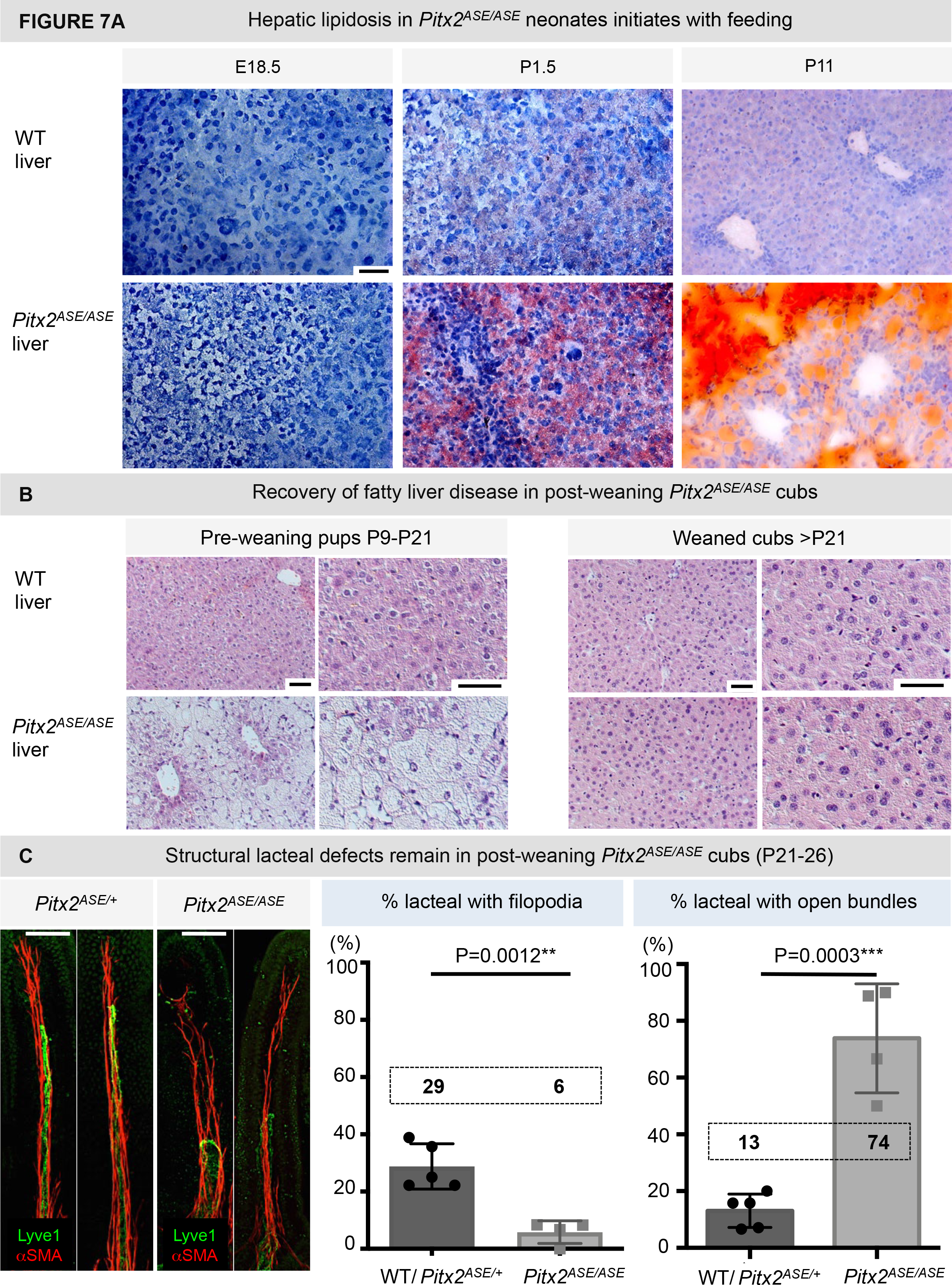
*Pitx2^ASE/ASE^* mice develop diet-induced fatty liver disease. (A) Hematoxylin (blue) and oil red O (red) staining of frozen liver sections from E18.5, P1.5, and P11 WT (top) and *Pitx2^ASE/ASE^* (bottom) mice. Note the enrichment of oil red O staining in postnatal, but not prenatal, *Pitx2^ASE/ASE^* livers. Scale bar, 50 μm. (B) Histopathology of WT (top) and *Pitx2^ASE/ASE^* (bottom) livers isolated from suckling pups (left) and weanlings (right). Note lipid vacuoles and loss of eosinophilic parenchymal stain in the liver of *Pitx2^ASE/ASE^* suckling pups, while lesions were resolved in *Pitx2^ASE/ASE^* weanlings. Images shown on the right side are selected high magnification fields of the left adjacent image. Scale bar, 50 μm. (C) Structural muscular-lacteal defects remain in post-weaning Pitx2*^ASE/ASE^* cubs. <u>Left</u>: Representative images of whole mount muscular-lacteal complexes from WT and *Pitx2^ASE/ASE^* cubs at P26. Note shorter lacteals (lyve-1, green) and more open muscular arrangement (αSMA, red) in *Pitx2^ASE/ASE^* but not WT weanlings. <u>Right</u>: Bar graphs showing quantifications of % lacteals with filopodia and with open muscular arrangement for each genotype. Note that the following mice were pooled: n=4 WT at P21 with n=1 *Pitx2^ASE/+^* at P26; *Pitx2^ASE/ASE^* group consisted of n=3 at P21 and n=1 at P26. Data displayed on Y-axes are represented as mean ± SEM. Lacteals with filopodia in WT weanlings = 28.81 ± 7.915 %, n = 5; lacteals with filopodia in *Pitx2^ASE/ASE^* weanlings = 5.833 ± 3.967%, n = 4. Difference between WT and *Pitx2^AS^*^E*/ASE*^ = -22.98 ± 4.375 %; P = 0.0012**. Lacteals with opened axial muscle in WT weanlings = 13.08 ± 2.637 %, n = 5; lacteals with opened axial muscle in *Pitx2^ASE/ASE^* weanlings = 73.85 ± 9.599 %, n = 4. Difference between WT and *Pitx2^ASE/ASE^* = 60.77 ± 8.945 %; P = 0.0003***. Black characters represent the mean of each group. Scale bar, 50 μm. See also Figure S7.

At P9-P11, oil red O staining was even more robust (Fig. 7A). Moreover, H&E analysis of affected pups at P9 revealed excessive hepatic lipid accumulation in 92% (n=11/12) of Pitx2*^ASE/ASE^* pups (Fig. S7B), with variable parenchymal extinction and multifocal subcapsular mineralization (Fig. 7B). Importantly, no significant changes in hepatic structure, hepatocyte mitochondria morphology, or extramedullary hematopoiesis were observed between Pitx2*^ASE/ASE^* and WT littermates (data not shown), ruling out primary liver impairment.

While the lipid content of murine maternal milk is ∼ 17-30% [77], the lipid content in standard mouse chow diet is 6-10 fold lower (∼ 3%). We noticed that hepatic lipidosis in *Pitx2^ASE/ASE^* suckling pups gradually resolved once these pups were weaned (P21) and transitioned to a dry pellet diet (Fig. 7B). Severe hepatic lipidosis in *Pitx2^ASE/ASE^* pups was seen in only 50% (n=2/4) of pups at P21 (compared to 92% at P9) and was further reduced to 25% (n=1/4) at P26, and 0% after P30 (n=0/2) (Fig. S7B). Importantly, the intestinal lymphatic lesions caused by *ASE* deletion did not resolve with age or weaning or a change of diet. This is a strong indication that adult mature mice remain susceptible to lipid accumulation dependent on their diet. These lesions included lacteal filopodia defects (Fig. 7C; WT, 28.81 ± 7.915%, n=5; Pitx2*^ASE/ASE^*, 5.833 ± 3.967%, n=4, p=0.0012), open versus closed axial smooth muscle (Fig. 7C; WT, 13.08 ± 2.637%, n=5; Pitx2*^ASE/ASE^*, 73.85 ± 9.599%, n=4, p= 0.0003), and severe axial smooth muscle hypoplasia (Fig. S3B). Collectively, these data suggest that high fat diet (in this case maternal milk) drives the pathogenesis of fatty liver in Pitx2*^ASE/ASE^* pups, and that Pitx2c/*ASE* protects against such changes by directing normal villous smooth muscle morphogenesis and the dependent process of gut lymphatic development.

## DISCUSSION

We have leveraged the *ASE* mouse model to characterize intestinal lymphatics and lipid transport function in the absence of the LR asymmetric transcription factor *Pitx2*. Our work demonstrates a new role for the LR axis in the development and physiology of intestinal lymphatics, linking gut lymphatic morphogenesis to the earliest LR symmetry-breaking events in the gut. We found that *Pitx2* functions during lacteal morphogenesis by regulating cellular redox in the adjacent smooth muscle lineage. We propose that lacteal-associated contractile SMCs are of blood vessel origin and arise during embryonic development by *ASE*-dependent remodeling of villous blood capillary-associated muscle progenitors to support dietary lipid transport soon after birth. *ASE* deletion, which leads to malformation of the SMC-lacteal complex results in abnormal lipid trafficking, which underscores the important reciprocal interplay between smooth muscle development and lacteal function in dietary lipid transport. Collectively, our studies highlight the critical integration of LR patterning signals that asymmetric viscera must execute from early axis specification through to mature organ function.

### *Pitx2* in intestinal smooth muscle development

The evolutionarily conserved process of LR asymmetry is a fundamental aspect of the body plan, errors in which form an important class of human birth defects. Whereas the intestine begins as a symmetrical midline tube, it later rotates and loops in a highly conserved, asymmetric pattern necessary for correct packing into the body cavity. Gut asymmetry initiates with a critical leftward tilt driven by molecular and cellular asymmetries within the left and right sides of the dorsal mesentery, which suspends the gut tube. These events are orchestrated by *Pitx2* expressed in all cells of the left dorsal mesentery [16]. To avoid strangulation of gut vessels during looping morphogenesis, *Pitx2*-dependent mechanisms directing initial gut rotation also pattern the emerging gut vasculature, linking the complex steps of vascularization with the morphogenesis of the parent organ [11, 78].

In addition to directing asymmetric gut development, the dorsal mesentery is the major source of smooth muscle progenitors of the gut vasculature [79, 80]. This raises the possibility that *Pitx2* may also regulate intestinal smooth muscle patterning. Using lineage tracing experiments to follow *Pitx2* (and *ASE*) daughter cells in the gut, we show that *Pitx2* descendants populate multiple intestinal smooth muscle layers, including the lacteal axial SMCs that drive lacteal contraction and efficient lipid transport in the postnatal period [38]. Interestingly, not all axial SMC bundles within the villus are descendants of *Pitx2* (Fig. 3A”A”’), suggesting an additional source of muscle progenitors within the dorsal mesentery and raising important questions about the overall functional heterogeneity among lacteal-associated axial SMCs. Whereas *Pitx2* is the primary determinant in the left dorsal mesentery, the T-box transcription factor *Tbx18* is one of the most highly differentially expressed genes on the right side, and its asymmetric expression within the dorsal mesentery correlates with cellular behavior [16, 18]. In the kidney, *Tbx18*-derived progenitor cells give rise to vascular SMCs that originate from stromal precursors, and stromal cell differentiation requires *Tbx18* function to form the renal vasculature [81, 82]. Therefore, Tbx18+ cells within the right dorsal mesentery may provide an additional source of axial SMCs, functionally distinct from those on the left. Similar heterogeneity has recently been demonstrated within the villous fibroblast compartment, highlighting the importance of the dynamic paracrine milieu of the lamina propria to the overall integrity and function of intestinal lymphatic vasculature [83].

In addition to populating axial SMCs, *Pitx2* daughter cells were also found in close proximity to the villous blood capillary plexus suggesting that they are blood capillary-associated immature mesenchymal precursors to the more mature muscle (Fig. 3A’A””). *Pitx2* daughter cells were also found in a subpopulation of the circular and longitudinal visceral smooth muscle layers within the muscularis externa (data not shown), which drives lymph propulsion through gut peristalsis, suggesting that *Pitx2* may regulate overall gut motility. Consistent with this idea, *Pitx2* serves to modulate the expression of several contractile proteins in the skeletal muscle including myosin heavy chain (MyHC) isoforms and the contractile regulatory proteins troponin I and troponin T [84]. Moreover, during myogenic development, *Pitx2* is downstream of Pax3/7 [85, 86] and upstream of the myogenic factor genes Myf5, Myf6, and MyoD [85, 86]. Mutations of *Pitx1*, *Pitx2*, and *Pitx3* result in developmental defects in humans, including muscle disorders such as Facioscapulohumeral Muscular Dystrophy [87], Axenfeld-Rieger syndrome [88], and Anterior Segment Mesenchymal Dysgenesis[89], respectively. Most of what is known about *Pitx2* in the molecular control of myogenesis concerns skeletal muscle, derived from paraxial mesoderm. Here, our work provides new insights into the function of *Pitx2* in the visceral smooth muscle from splanchnic mesoderm and suggest *Pitx2*-mediated mechanisms are involved in intestinal SMC morphogenesis. We speculate that the regulation of muscle biology is a common property of Pitx transcription factors at their distinct spatiotemporal sites of expression.

### *Pitx2*-dependent formation of the lacteal-associated axial SMC complex

Although recent studies have revealed novel insights into the formation of the muscularis externa [90], the origins and mechanisms of lacteal-associated axial SMC formation remain unexplored. Our spatiotemporal characterization of villous muscular formation demonstrate that the cellular origin of axial SMCs does not appear to be a continuous extension from the submucosal lymphatic vasculature, as these vessels are devoid of muscle coverage [29]. Instead, our data suggest that lacteal-associated SMCs arise from villous blood capillary-associated star-like muscle progenitors in a process regulated by *ASE*. In the adult blood vessels, differentiated muscle cells proliferate at extremely low rates, exhibit low synthetic activity, and express a unique repertoire of contractile proteins [49, 91]. Consistent with axial SMCs resembling a more differentiated muscle cell type, we show that they exhibit a low rate of proliferation, are strongly αSMA-positive, and express high levels of Myh11. In contrast, αSMA+ star cells are proliferative and show weak immunoreactivity for αSMA and Myh11. Thus, our data suggest that blood capillary-associated star cells likely represent less differentiated smooth muscle progenitor phenotype and lie upstream of the more mature bundles of axial SMCs. Of note, a small number of αSMA+ star cells remains in WT villi at P9 (Fig. S4D). This raises the possibility that these cells may contribute to further postnatal axial muscular development or regeneration during adult life. It is also conceivable that many more star-like progenitors remain in the villus but have lost αSMA expression akin to SMC phenotypic switching [47, 48]. These cells would be undetectable by smooth muscle markers postnatally.

The timely transition from αSMA+ star cells to elongated spindle-like αSMA++ cell types coincided with the appearance of lacteals (E18.5) implying bidirectional crosstalk between lacteal development and the onset of axial SMC remodeling. Postnatal lacteals were always accompanied by the adjacent axial SMCs and our analysis of WT and *Pitx2^ASE/ASE^* intestines failed to detect lacteals without their muscular neighbors. However, we did observe instances of axial SMCs in villi with missing lacteals in *Pitx2^ASE/ASE^* intestines (Fig. 3B). Importantly, axial SMCs that formed in the absence of lacteals were always mispatterned with aberrant branches interconnecting the bundles (Fig. 3B). Together, these data suggest that axial SMCs are necessary for lacteal formation. And while lacteals are dispensable for the formation of axial SMCs, they are required for proper muscle bundle organization. Thus, *ASE* deletion, which leads to malformation of the muscular-lacteal complex, reveals the important reciprocal interplay between lacteal development and axial SMC remodeling during the pre- to postnatal transition.

### *Pitx2*-mediated changes in ROS levels in the intestinal muscle

Previous studies have shown that oxidative stress inhibits mesenteric lymphatic vessel contractility and reduces lymph flow by affecting lymphatic smooth muscles in anesthetized and aging rats [92, 93]. Consistent with the role of ROS in lymphatic function, our studies suggest that elevated levels of oxidative stress caused the muscular-lacteal impairment in *Pitx2^ASE/ASE^* intestines pointing to a crucial role for *Pitx2* in the control of ROS during early development. During normal muscle development, the transition of proliferative muscular progenitor into postmitotic contractile mature muscle involves a glycolytic to oxidative metabolic switch, leading to excessive production of ROS [94, 95]. Whereas this can damage cells, elevated ROS are crucial to normal physiological responses such as differentiation [96, 97]. Therefore, a network of genes counteracting elevated ROS is critical to mitigate cell damage. We propose that *Pitx2* is one of those genes in the intestine. First, Pitx2*^ASE/ASE^* neonates had significantly more intestinal protein carbonyl content in comparison to their WT or Pitx2*^ASE/+^* heterozygous littermates. Second, they had the most carbonylated αSMA of all genotypes tested, demonstrating increased irreversible actin oxidation in the absence of *ASE*. Based on our RNA-seq data, these effects are likely driven by the cumulative effects of disturbed redox homeostasis-related genes downstream of Pitx2, including Gclc [50–52].

Actin carbonylation (indicative of both F- and G-actin oxidation) is indicative of severe functional impairment of the actin cytoskeleton [61]. Importantly, due to the irreversible nature of protein carbonylation, contractile proteins exposed to early oxidative stress are damaged permanently. In other words, an early disruption in actin dynamics may have repercussions that persist at later stages. In that regard, we observed fragmented patterns of αSMA arrangement in *Pitx2^ASE/ASE^* axial SMCs at P21 (Fig. S5B) and profound villus smooth muscle hypoplasia at P26 (Fig. S3B). Whereas we were unable to characterize further muscle morphogenesis due to the early postnatal lethality of *Pitx2^ASE/ASE^* mice, our data suggest that Pitx2-mediated control of ROS is a critical early step to maintaining an organized and dynamic actin cytoskeleton during both organogenesis and subsequent contractile muscle function later in life. Consistent with this notion, recent studies found an essential role for Pitx2 and Pitx3 in the management of ROS in differentiating myoblast and satellite stem cells [98]. Furthermore, in the neonatal heart, *Pitx2c* was shown to inhibit ROS after cardiac injury to promote heart repair by activating the antioxidant response pathway and ROS scavenger genes [99]. Collectively, these data implicate Pitx2 as a core rheostat of redox status during muscle development and regeneration and necessitate future studies targeted to defining the specific roles for *Pitx2c* in smooth muscle regeneration after intestinal damage.

### *Pitx2* regulates lacteal development and function in a non-cell autonomous manner

It is well established that lymphatic homeostasis and function is regulated in part in a non-cell autonomous manner by the adjacent microenvironment [35, 37, 83, 100, 101]. Because ROS can act on surrounding cells with [102, 103] or without [104] diffusion, our data support this model and raise the possibility that ROS-generating axial SMCs can influence cell behavior of the adjacent lacteal including the formation of lacteal filopodia extensions. Filopodia are formed by linear polymerization of G-actin-ATP at their barbed–ends, and the lengthening and retraction of filopodia actin bundles are regulated by ROS [105–107]. During intestinal lymphatic development, lacteal filopodia are crucial for lacteal sprouting, extension, and regeneration throughout adulthood, ensuring continuous lacteal function for transport of dietary lipids [33, 35]. During adult lacteal regeneration, filopodia-mediated lacteal migration is regulated by the Notch ligand delta-like 4 (DLL4), downstream of VEGF-C [35]. Genetic inactivation of DLL4 in lymphatic endothelial cells [35] or postnatal deletion of VEGF-C [37] leads to lacteal regression and impaired dietary fat uptake. Moreover, interaction between NRP2, a transmembrane co-receptor for VEGF-C, and VEGFR-3 mediates proper lacteal sprouting in response to VEGF-C [33]. Interestingly, we have previously identified three highly conserved *Pitx2* binding regions upstream of *VegfC* transcriptional start sites by ChIP-seq analysis and ChIP-qPCR of embryonic intestines and detected reduced *VegfC* expression in *Pitx2^ASE/ASE^* embryos as early as in E10.75 (data not shown). While VEGF-C has been implicated in multiple crucial aspects of lymphangiogenesis including intestinal lymphatics [37], how this signaling is integrated in the context of a developing organ remains unknown and our preliminary data suggest that Pitx2 may provide this context. Collectively, elevated levels of ROS in *Pitx2^ASE/ASE^* intestines and reduced VEGF-C- VEGFR3 signaling in the absence of *ASE* may have damaging effects on filopodia-mediated lacteal extension, resulting in shorter lacteals and impaired dietary fat uptake and/or trafficking.

### *Pitx2^ASE/ASE^* mice develop diet-induced fatty liver disease

In viviparous mammals, a functional intestinal lymphatic system is vital for rapid processing of fat-enriched milk at birth, and in lipid absorption through adulthood. Dietary lipids (mostly long-chain fatty acids), cholesterol, and fat-soluble vitamins use lacteals as the major transport route, whereby lipids packaged into lipoprotein particles (chylomicrons) drain into larger mesenteric collecting lymphatic vessels, and ultimately into the systemic blood circulation. In contrast, non-fatty nutrients and shorter chain fatty acids enter the portal vein directly for delivery to the liver [9, 30–32]. This portal transport is insignificant compared with the chylomicron pathway, but can become active during disease where disrupted lymphatic clearance impairs the normal route of lipid transport [71-73, 108, 109]. Importantly, whereas lymphatic-dependent transport distributes dietary lipids to peripheral tissues for direct energy consumption, the lymphatic-independent lipid trafficking directs lipids exclusively to the liver for immediate catabolism by hepatocytes. This greatly reduces the lipid accessibility to peripheral tissues and may be the cause of growth retardation seen in the *Pitx2^ASE/ASE^* mice.

In our model, the lymphatic-dependent lipid transport is dominant in WT neonates, directing dietary long-chain fatty acids into the systemic lymphatic circulation. In contrast, muscular-lacteal development and contractions in the absence of *ASE* are insufficient to support lymphatic-dependent lipid transport. As a consequence, lymphatic-independent lipid trafficking is activated to cargo lipids from the intestine, leading to excessive hepatic lipid accumulation and fatty liver in neonatal *Pitx2^ASE/ASE^* mice. Interestingly, the fatty lesions were reversible when *Pitx2^ASE/ASE^* weanlings transitioned to a low fat diet. This suggests that the duration of fat enriched diet is critical to the prognosis of fatty liver disease, where non-alcoholic fatty liver disease (NAFLD) may be resolved if early diet adjustments are made, while extended high fat diet may lead to non-alcoholic steatohepatitis (NASH).

How lipids are shunted to the portal circulation rather than the lymph in the *Pitx2^ASE/ASE^* intestine is a critical and fascinating question. A majority of liver perfusion derives from the portal vein and any portal venous cargo strongly impacts liver function (as in first-pass drug metabolism). Dietary long-chain fatty acids along with other nutrients, including shorter chain fatty acids, carbohydrates, and amino acids are absorbed by enterocytes, a function that is broadly zonated along the villus axis: genes associated with chylomicron packaging and lipid transport functions are most highly expressed in the tip-most enterocytes whereas amino acid and carbohydrate transporters are enriched within the middle of intestinal villi [110]. These fascinating data suggest a chylomicron concentration gradient along the villus axis, consistent with prior tEM analysis [111]. Accordingly, taller lacteals contain more chylomicrons than shorter lacteals. Moreover, lacteals are positioned further away from the enterocytes than the venous capillaries, giving priority for small nutrients entry into the portal circulation. Thus, we postulate that shorter lacteals in *Pitx2^ASE/ASE^* mice have a net deficit in chylomicron entry overall, further aggravating lipid transport insufficiency. Fatty liver disease is currently the world’s most common chronic liver disease [112]. Here, our studies link this disease to altered gut lymphatic development and transport of fat in the intestine, pointing to a novel role for the Pitx2-driven organ laterality in the regulation of the gut-liver axis. Understanding the gut-liver axis in relation to lipid trafficking through portal versus lymphatic vasculature is critically needed, as it will lead to future discoveries of alternative regulators of fat absorption that may be clinically targeted under disease conditions. Taken together, our findings unravel a previously unknown role for the LR *Pitx2* gene during muscular-lacteal morphogenesis and reinforce the importance of the reciprocal interplay between intestinal muscle development and lacteal function in dietary lipid transport.

## ACKNOWLEDGMENTS

We thank all the Kurpios lab members and Dr. D. Gludish for reading the manuscript and excellent suggestions. We thank Drs. H. Hamada, J. F. Martin, and T. Tumbar for reagents described above. We thank T. Bargar and N. Conoan of the Electron Microscopy Core Facility at the University of Nebraska Medical Center for technical assistance, supported by state funds from the Nebraska Research Initiative and the University of Nebraska Foundation, and institutionally by the Office of the Vice Chancellor for Research. We are grateful to Dr. B. Dixon, Dr. T. Cassis, and A. VanDeMark for help with BODIPY feeding of newborn mice, Drs. J. Dela Cruz and R. Williams for imaging expertise, A. Sulpizio for help with lipid quantifications, and Drs. T. Stokol, E. L. Behling-Kelly, T. L. Southard, and S. Center for clinical discussions. We thank Dr. Jen Grenier at the Transcriptional Regulation and Expression Facility and the Biotechnology Resource Center (BRC) of Cornell Institute of Biotechnology for RNA-seq analysis and R. Munroe and C. Abratte of the Cornell Stem Cell and Transgenic Core Facility. We are grateful to B. Laslow, R. Slater, and C. Westmiller for technical assistance. Imaging data was acquired through Cornell BRC Imaging Facility, with NYSTEM (C029155) and NIH (S10OD018516) funding for the shared Zeiss LSM880 confocal/multiphoton microscope. NIH-funded Comparative Medicine Training Program T32OD011000 (S.H.), AHA-funded 17SDG33400141 (G.T.), and NIDDK R01 DK107634 and DK092776 (N.A.K.) supported this work.

## AUTHOR CONTRIBUTIONS

Conceptualization, S.H., A.M., and N.A.K.; Methodology, S.H., A.M., C.S.S., and N.A.K.; Investigation, S.H., A.M., I.F.E., J.C., N.R.S., G.H.B., A.K.T., and G.T.; Writing – Original Draft, S.H., Writing – Review & Editing, S.H. and N.A.K.; Funding Acquisition, S.H. and N.A.K.; Supervision, G.E.D., C.S.S., and N.A.K.

## DECLARATION OF INTERESTS

The authors declare no competing interests

## ACCESSION NUMBERS

The RNA-sequencing data have been deposited under accession number GEO: GSE160677

## Methods

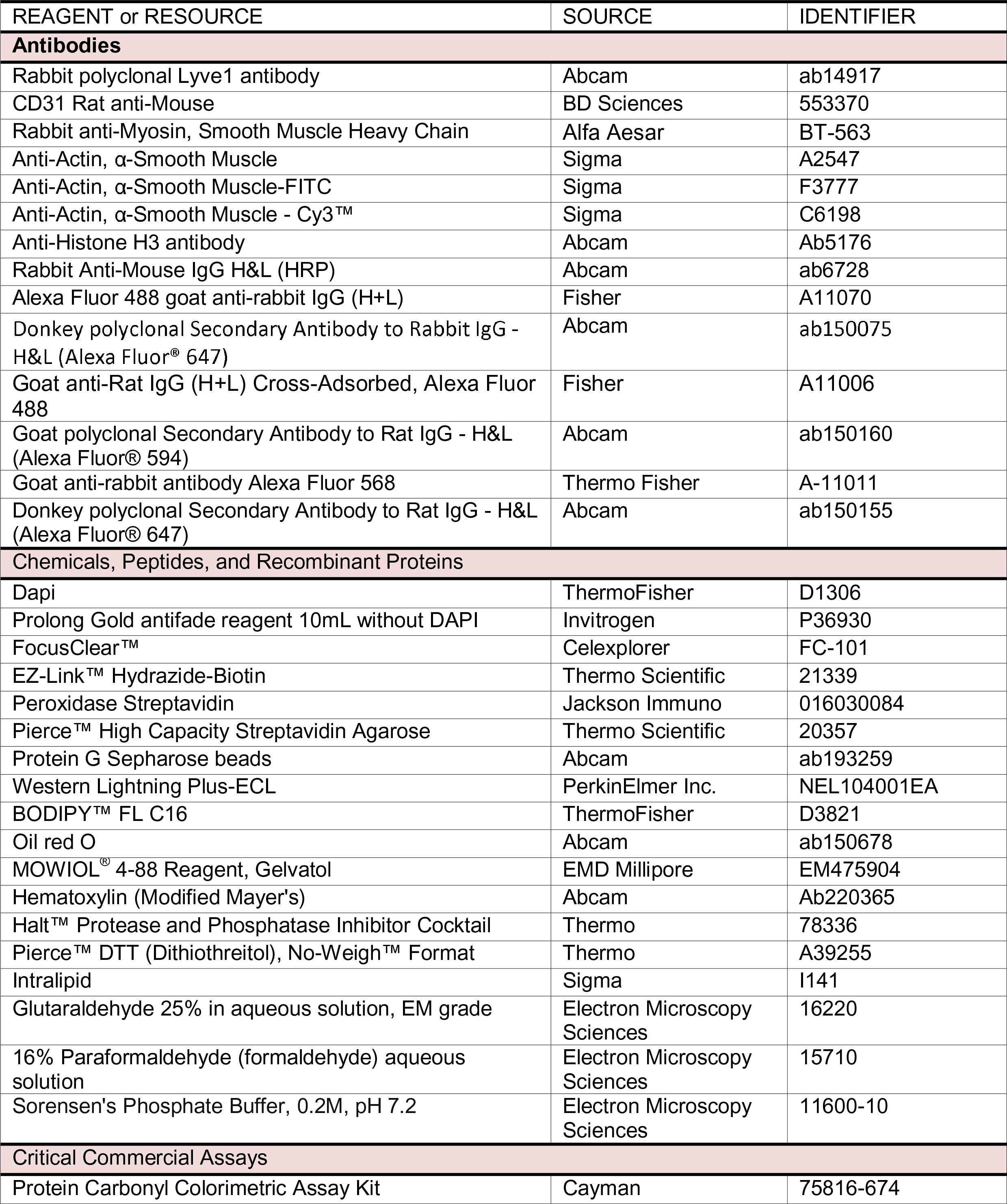
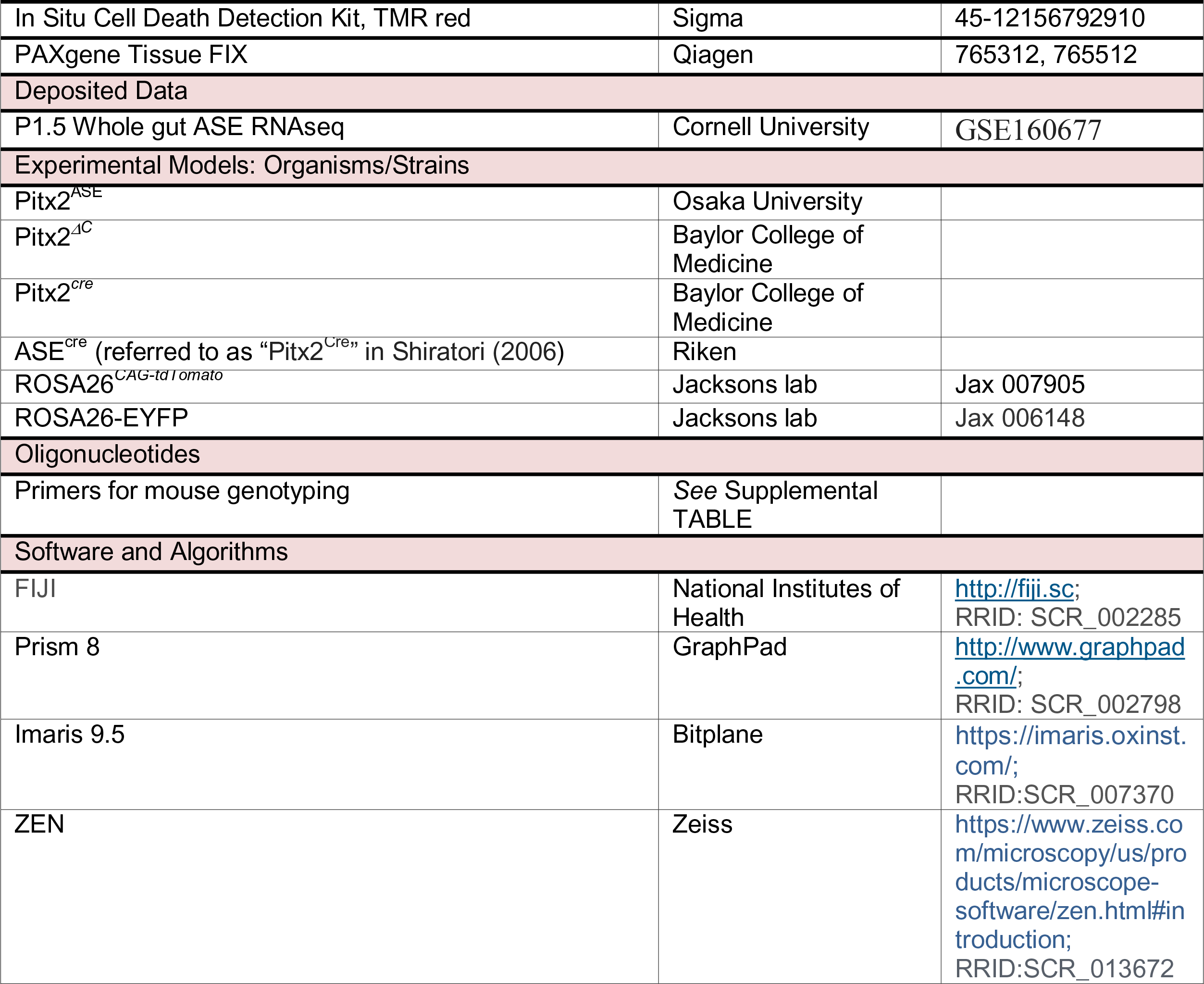
Key Resources Table.

### Resource Availability

#### Lead contact

Further information and requests for resources and reagents should be directed to and will be fulfilled by the Lead Contact, Natasza A. Kurpios.

### Materials Availability

This study did not generate new unique reagents.

### Data and Code Availability

The RNAseq database generated during this study is available at GEO: GSE160677

Experimental Model and Subject Details

### Mice

All experiments adhered to guidelines of the Institutional Animal Care and Use Committee of Cornell University, under the Animal Welfare Assurance on file with the Office of Laboratory Animal Welfare. Pitx2^ASE/ASE^, Pitx2*^ΔC^*, Pitx2^cre^, ASE^cre^, ROSA26*^CAG-tdTomato^*, and ROSA26-EYFP mice were previously described [22, 23, 28, 113, 114]. All strains of mice in these studies were maintained in a specified pathogen-free barrier facility under a 12-hour light cycle with free access to standard chow pellets and water unless specified. All mice used for mating were between 2-6 months old. For timed pregnancies, dams and studs were housed separately until the scheduled mating. Time mated breeding pairs were separated upon the morning when vaginal plugs were found and staged as E0.5. Postnatal stages are defined upon birth as postnatal day 0 (P0). Each mouse used was numbered by toe clip and genotyped by PCR amplification at the age between 7-10 days. Embryos and pups under 9 days were genotyped after isolation. For full list of genotyping primers, see Supplemental Table. Each group of comparison contained at least three mice from representative genotypes at the specified developmental stages. Mice of different genotypes were randomly chosen and sorted into different procedures and measurements. When possible, littermates of different genotypes were sorted into the same procedure and measurements as pairs.

## Method Details

### Mouse tissue collection, fixation, processing

For sectioning, tissues were dissected in ice-cold 1X PBS, immediately followed by PBS washes and fixed overnight at 4°C in 4% paraformaldehyde (PFA)/PBS, ice-cold 100% MeOH, or PAXgene® fixative. For tissues fixed in PFA, tissues were thoroughly washed with PBS, then preserved in 4°C until use. Tissues fixed in MeOH were stored in -20°C until further use. Tissues fixed with PAXgene® fixative were processed following manufacturer’s instructions. For intestinal tissue that proceeded to frozen sections, tissues were dehydrated in gradient sucrose/PBS solution, then embedded in Tissue-Tek* OCT. Compound and stored in -80°C until sectioning. Hepatic tissue for frozen sections following oil red O stain was embedded in 100% OCT immediately after dissection. Tissues intended for histology analyses were processed with standard paraffin embedding protocol in the Animal Health Diagnosis Center, Cornell University College of Veterinary Medicine. All frozen sections were 15 μm thick; paraffin sections were 5 μm thick.

### Section and whole mount immunofluorescence

Embryos were processed for immunofluorescence (IF) on sections or whole mount (tissue slice). For section IF, frozen sections were permeabilized with 0.1% TritonX-100/PBS for 45 min, blocked in 3% BSA in PBS for 3 hours at room temperature, and incubated with 100X diluted primary antibodies/blocking overnight at room temperature. 500X diluted secondary antibodies were co-incubated with 1000x diluted DAPI for nuclear labeling at room temperature for 1 hour. Samples were mounted in ProLong^TM^ Gold Antifade Mountant (P36930, Thermo Fisher) and preserved in room temperature until imaging. For whole mount IF staining, the protocol was adapted from [43]. In brief, tissue was blocked in BSA, serum, and 0.3% TritonX-100 in PBS in 4°C for 3 hours, then incubated with primary antibodies overnight at 4°C. Tissues were thoroughly washed in 0.3% TritonX-100/PBS for 5 hours with hourly buffer changes, then incubated with secondary antibody overnight at 4°C. Tissue was thoroughly washed in 0.3% TritonX-100/PBS for 5 hours with buffer changes every half hour, then fixed in 4% PFA/PBS for two overnights at 4°C. For confocal imaging, tissue was washed with PBS and properly sliced to 100-200 μm with spring scissors before mounted in ProLong^TM^ Gold Antifade Mountant. Primary antibodies are listed below: GFP (ab290, Abcam), CD31 (PECAM-1, 553371, BD Bioscience), Lyve-1 (Ab14917, Abcam), αSMA (F3777, C6198, A2547, Sigma-Aldrich), and Myh11 (BT-563, Alfa Aesar). Secondary antibodies used are listed below: AlexaFluor anti-goat, anti-mouse or anti-rabbit 568, and anti-rabbit 488 (source).

### Liver oil red O stain

Oil red O stain was performed to detect liver neutral lipids as previously described [76]. In brief, tissue sections were processed as described in the previous section. Slides with paired tissues from WT and *Pitx2^ASE/ASE^* littermates collected on the same slide were equivalate to room temperature from -80°C when stored, following oil red O and hematoxylin incubation. Slides were rinsed in water before mounted in MOWIOL^®^ 4-88 Reagent (EM475904, EMD Millipore).

### *In vivo* lipid tracer feeding assay

BODIPY^TM^FL C16 (Thermo Fisher D3821) was added to a warmed Intralipid (20% emulsion, Sigma I141) solubilizing agent to make final concentration at 0.4μg/μl. 50 μL of 37°C reconstituted Bodipy^TM^/Intralipid solution was fed to each pup through 24-gauge reusable feeding needles (Fine Science Tools 18061-24). To maintain proper body temperature and hydration, the pups were left with the dam for 5 hours (milk *ad lib*) before imaging and tissue collection. Images were taken under a Zeiss dissection microscope immediately after sacrifice by decapitation. Images were taken from liver tissue flash frozen in liquid nitrogen, then cryo-sectioned at 20 μm and mounted in ProLong^TM^ Gold Antifade Mountant.

### Tissue carbonyl assay detection

Protein Carbonyl Colorimetric Assay Kit (Cayman 10005020) was adapted for intestinal protein carbonyl measurement. In brief, small intestines were collected from animals euthanized by hypothermia followed by PBS perfusion. Intestine contents were removed by gentle swirling in ice-cold PBS after exposing the interior. Liquids were removed as much as possible before flash freezing the tissue in liquid nitrogen. 150 mg of tissue was homogenized in 50 mM phosphate buffer pH 6.7 then centrifuged at 10,000 x g for 15 min at 4°C. Supernatants were collected and incubated with 1% streptomycin sulfate in 50mM potassium phosphate pH 7.2 at room temperature following centrifugation at 6,000 x g for 10 min at 4°C to remove excessive nucleic acids. Supernatants were collected and stored in -80°C until further use. The tissue carbonyl assay was then performed and analyzed following manual instruction.

### Carbonylated total protein and smooth muscle actin-alpha measurement

EZ-link hydrazine-biotin was used to label protein carbonyl as previously described [115, 116]. Small intestines were collected in ice-cold PBS following removal of intestinal contents. Tissues were stored in -80°C until further processed. In brief, tissues were homogenized in ice-cold homogenous buffer (pH5.5-100mM sodium acetate, 20mM NaCl, 0.1mM EDTA with protease and phosphatase inhibitor cocktail). Supernatant was incubated with 2% SDS in homogenous buffer in 65°C for 5 min after centrifugation at 1200 rpm for 10 min in 4 °C. Samples were further centrifuged at 13,000 g for 1 hour in 4°C. Supernatant was collected for protein concentration measurement and 100-150 μg of protein samples were then incubated with EZ-link hydrazine-biotin at 125uM for 2 hours at room temperature with constant mixing. Hydrazine-labeled samples were then dialyzed into PBS for streptavidin pulldown or αSMA IP. Specific bands of hydrazine-labeled proteins and αSMA were detected by western blot as previously described (REF or above).

### Transmission electron microscopy

Tissues were isolated in ice-cold PBS and gently agitated to remove blood. Liver tissues were cut into approximately 1mm size cubes and fixed in 2% Glutaraldehyde, 2% Paraformaldehyde in 0.1M Sorensen’s Phosphate Buffer overnight in 4°C then washed in ice-cold PBS 3 times. Tissues were then post fixed in 1% Osmium Tetroxide aqueous for 1 hour before they were dehydrated in graded ethanol series following Propylene Oxide washes and immersed in Araldite/Propylene Oxide 1:1 solution overnight. The samples were then soaked in fresh Araldite for 4 hours before final embedding and heated to polymerize in silicon rubber molds. Thin sections (60-90nm) were cut on a Leica EM UC6 Ultramicrotome. Sections were collected onto 200 mesh copper grids and stained with 2% Uranyl Acetate and Reynold’s Lead Citrate before examined on the FEI Tecnai G2 Spirit Transmission Electron Microscope operated at 80 Kv.

### RNA-sequencing

4 WT and 3 *Pitx2^ASE/ASE^* P1.5 small intestines were included in the analysis. Total RNA was isolated with Trizol, with an extra chloroform extraction to remove residual phenol and addition of glyco-blue as a carrier to promote RNA precipitation. RNA concentration was determined on a Nanodrop. TruSeq-barcoded RNA libraries were generated with the NEBNext Ultra II Directional RNA Library Prep Kit (New England Biolabs) with polyA+ enrichment using 1 μg total RNA as input. The libraries were sequenced on a NextSeq500 with single-end 85nt reads. Illumina pipeline software v1.8 was used for base calling. Trim_galore was used to trim and filter reads (--nextseq 20 --length 50). STAR was used to map reads to the Mus musculus mm10 reference using Ensembl gene annotations and output raw counts (--quantMode GeneCounts). Differential gene expression was defined by false discovery rate (FDR, adjusted p value for multiple tests) <= 0.05 from raw counts between WT and *Pitx2^ASE/ASE^*.

## Quantification and Statistical analysis

### Imaging and quantifications

Microscopic images were acquired using a Zeiss Observer.Z1/Apotome. Confocal images were taken on Zeiss LSM880 confocal microscopes. To capture the entire lacteal and villus in whole mount immunofluorescent stain, multiple adjacent fields of images were taken under a 40X lens then stitched on Zen before exporting for further analyses. Imaris 9.5 was used for image processing, filtering, background subtraction, confocal image stacking, and quantifications. Measurements (lacteal length, filopodia counts etc.) were done on Imaris 9.5 or Fiji. Z stack images were all analyzed on Imaris 9.5.

### Statistics

Statistical analyses were performed in GraphPad Prism 8 (La Jolla, CA). Comparison between different genotypes were tested by unpaired, two tailed Student’s t test. All data were first analyzed as mean values of measurements (lacteal length, filopodia number, etc.) from multiple villi in multiple fields across the proximal small intestine for each animal. Lacteal filopodium was defined as cell protrusion longer than or equal to 6μm. The mean values from all animals were then plotted in groups based on genotype and developmental stages. Almost all data were expressed as mean of the means from each mouse ± standard error of mean (SEM).

### Supplemental Table

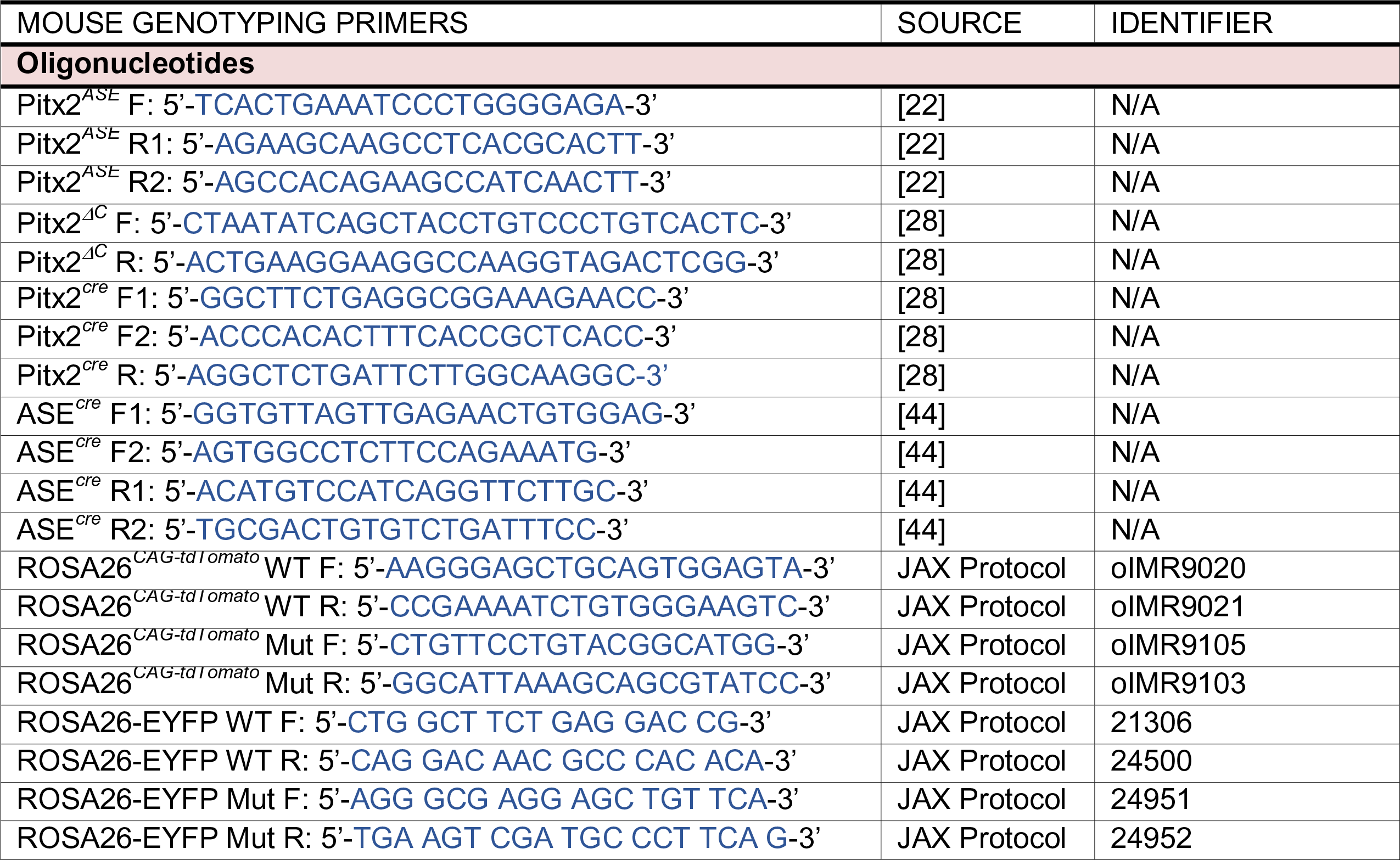

## SUPPLEMENTAL FIGURE LEGENDS

**Figure 8.**
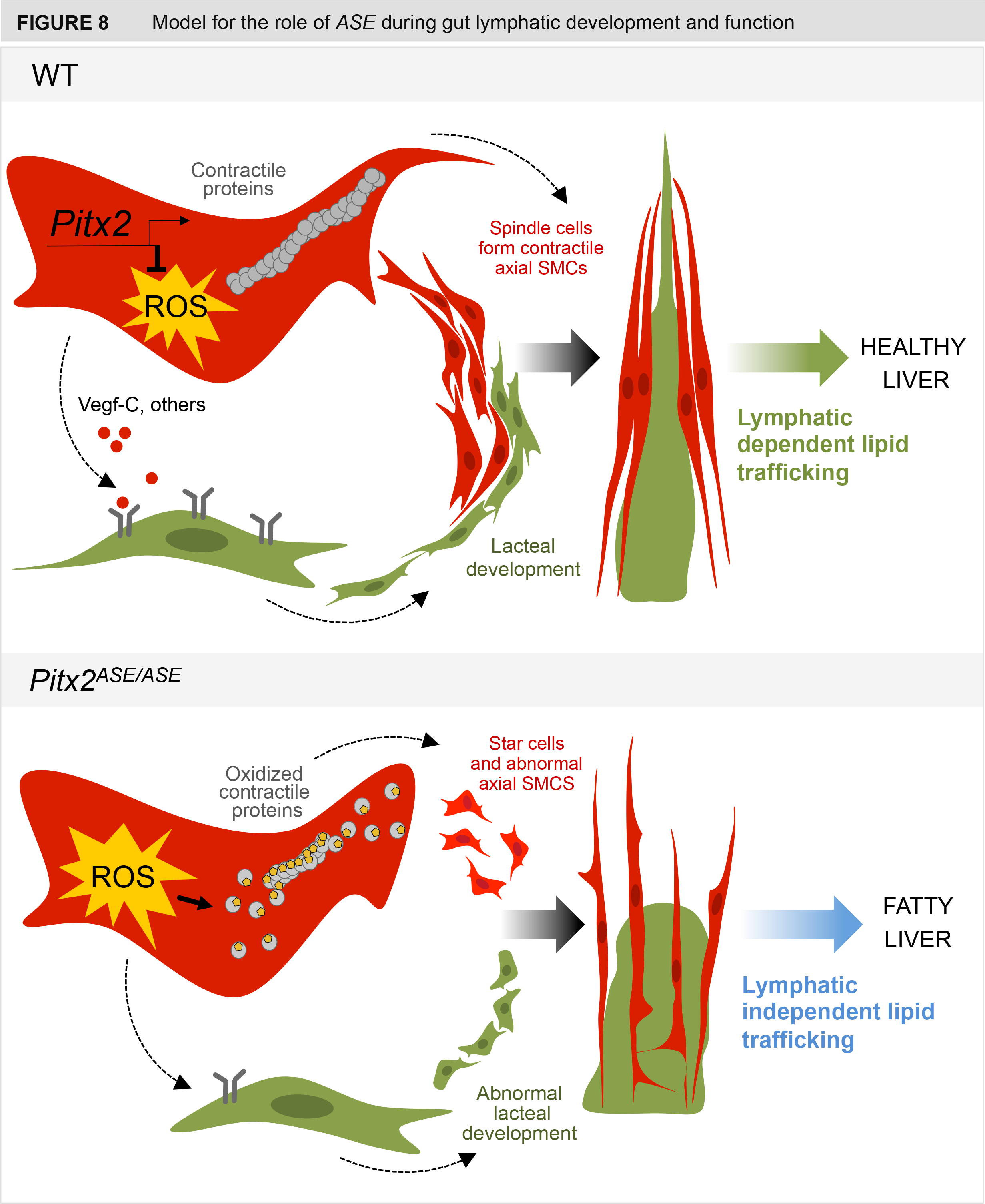
Proposed model for *Pitx2* function in gut lymphatics. <u>WT</u>: *Pitx2* (1) maintains redox balance during the development of villous SMCs, (2) supports SMC-dependent lacteal development, and (3) regulates lymphatic-dependent dietary lipid transport driven by villous SMCs. <u>Pitx2</u>*<u>^ASE/ASE^</u>*: sublethal *Pitx2* loss leads to (1) increased oxidative stress and protein damage in villous SMCs, therefore (2) compromises SMC-dependent lacteal development and (3) lymphatic-dependent lipid transport, while (4) potentially increases oxidative stress in adjacent LECs. This leads to activation of lymphatic-independent lipid transport, where excessive gut-liver lipid entry results in fatty liver disease.

**Figure S1.**
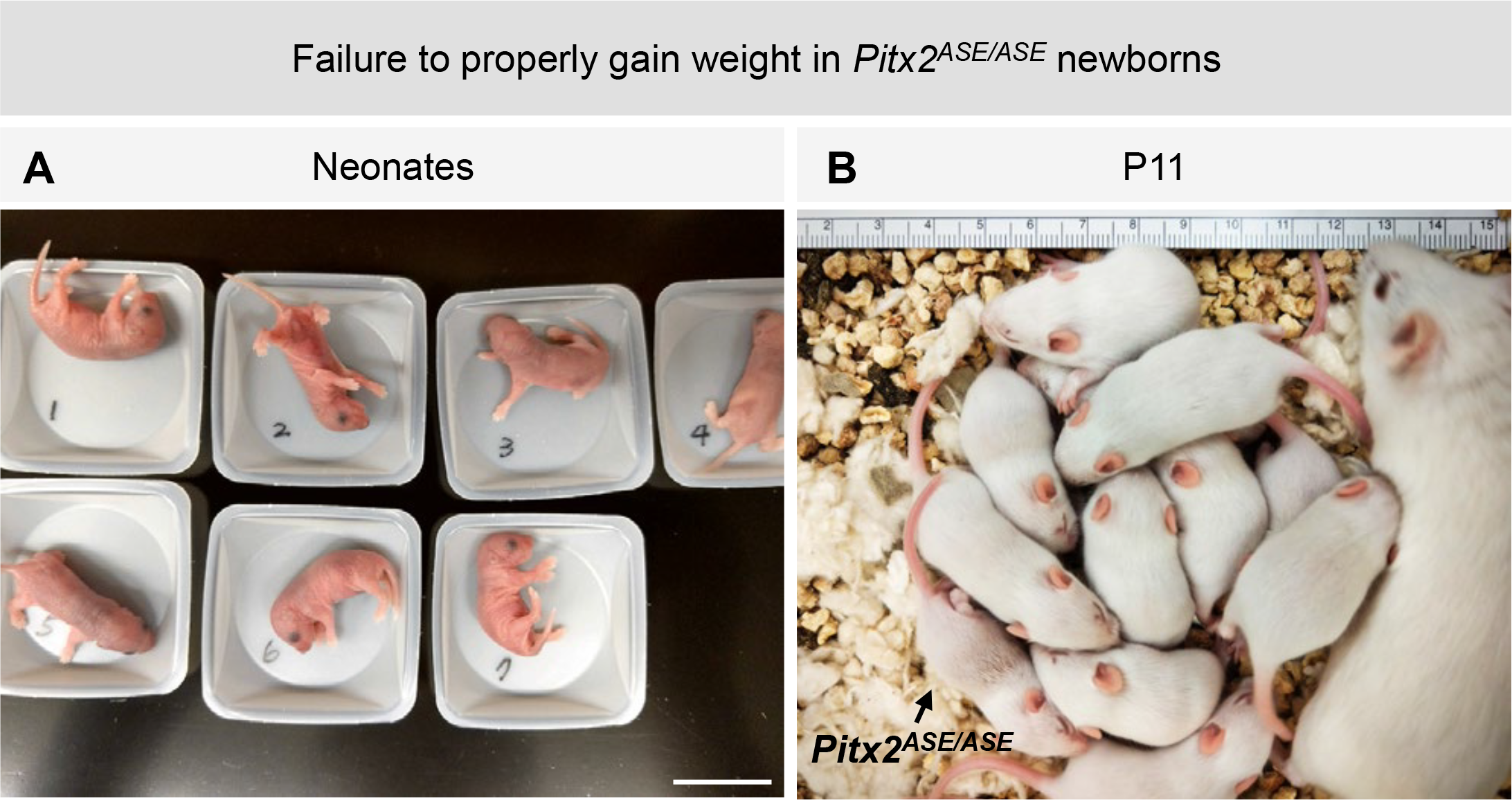
*Pitx2^ASE/ASE^* mice fail to properly gain weight (related to Figure 1) (A) Representative image showing a litter of newborn pups from a cross between two Pitx2*^ASE/+^* parents. Scale bar, 2cm. (B) Pitx2*^ASE/ASE^* pup is smaller than littermates (P11), but not excluded from the nursing litter. Note the Pitx2*^ASE/ASE^* runt in lower left corner.

**Figure S2.**
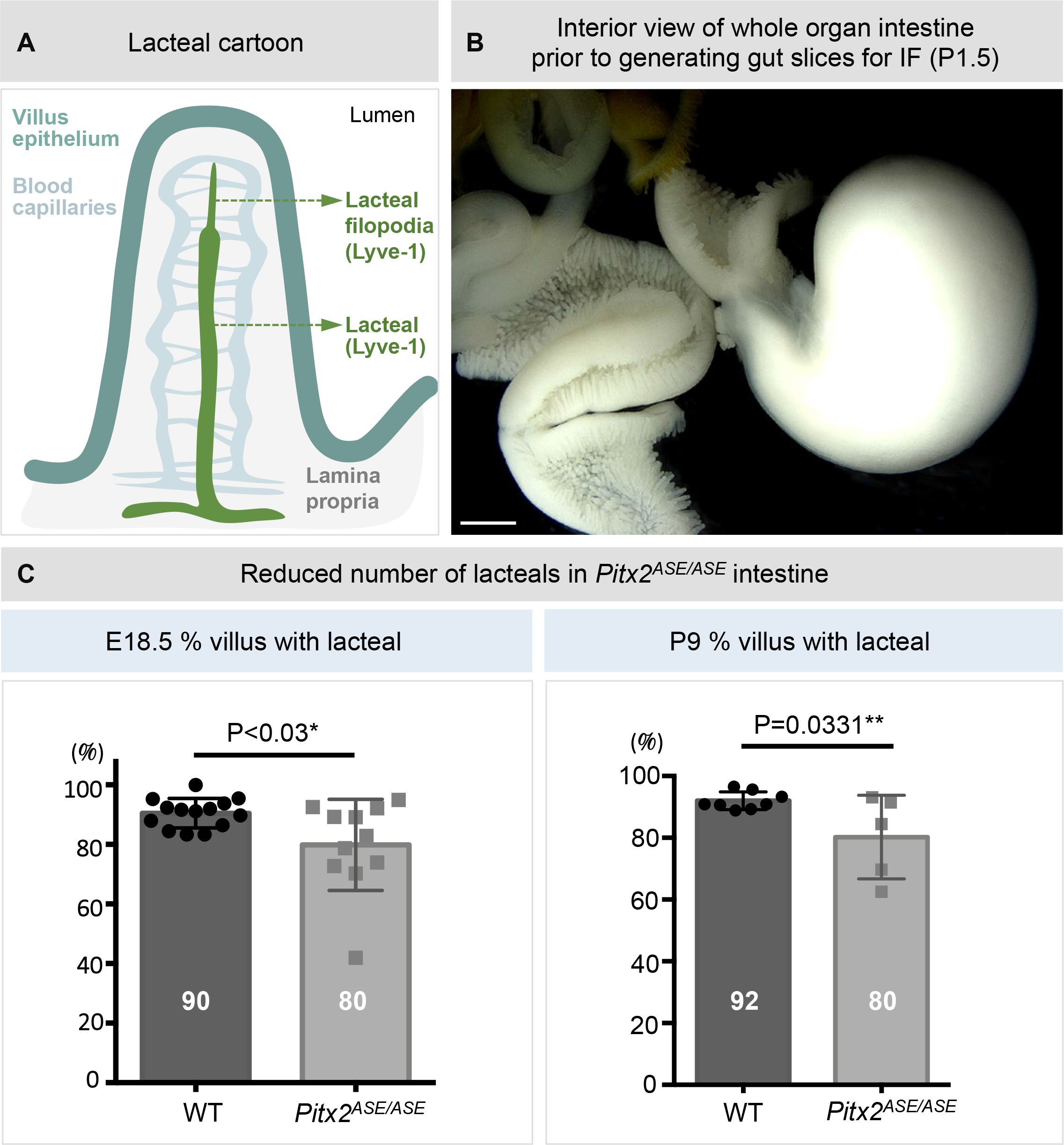
Whole mount analysis of small intestinal lacteal (related to Figure 2) (A) Cartoon showing the anatomical structure of lacteal and lacteal-filopodium. Green: lacteal endothelial cells; light blue: blood capillaries; teal: villus epithelium; gray: lamina propria. (B) Whole mount digestive tract at P1.5 with interior of the duodenum and jejunum exposed and intestinal content removed. Subsequent gut slices were generated for IF. Scale bar, 1000 μm. (C) Reduced number of villus with lacteals in pre- (left) and post- (right) natal Pitx2*^ASE/ASE^* by comparison to their WT littermates. Data displayed on Y-axes are represented as mean ± SEM % of villi with lacteal in each mouse. E18.5 WT = 90.49 ± 1.32 %, n = 14; E18.5 *Pitx2^ASE/ASE^* = 79.87 ± 4.617 %, n = 11. Difference between means = -10.62 ± 4.335 %. P = 0.0223*. P9 WT = 92.02 ± 1.010 %, n = 8; P9 *Pitx2^ASE/ASE^* = 80.24 ± 6.060 %, n = 5. Difference between WT and *Pitx2^ASE/ASE^* = -11.78 ± 4.836 %. P = 0.0331*

**Figure S3.**
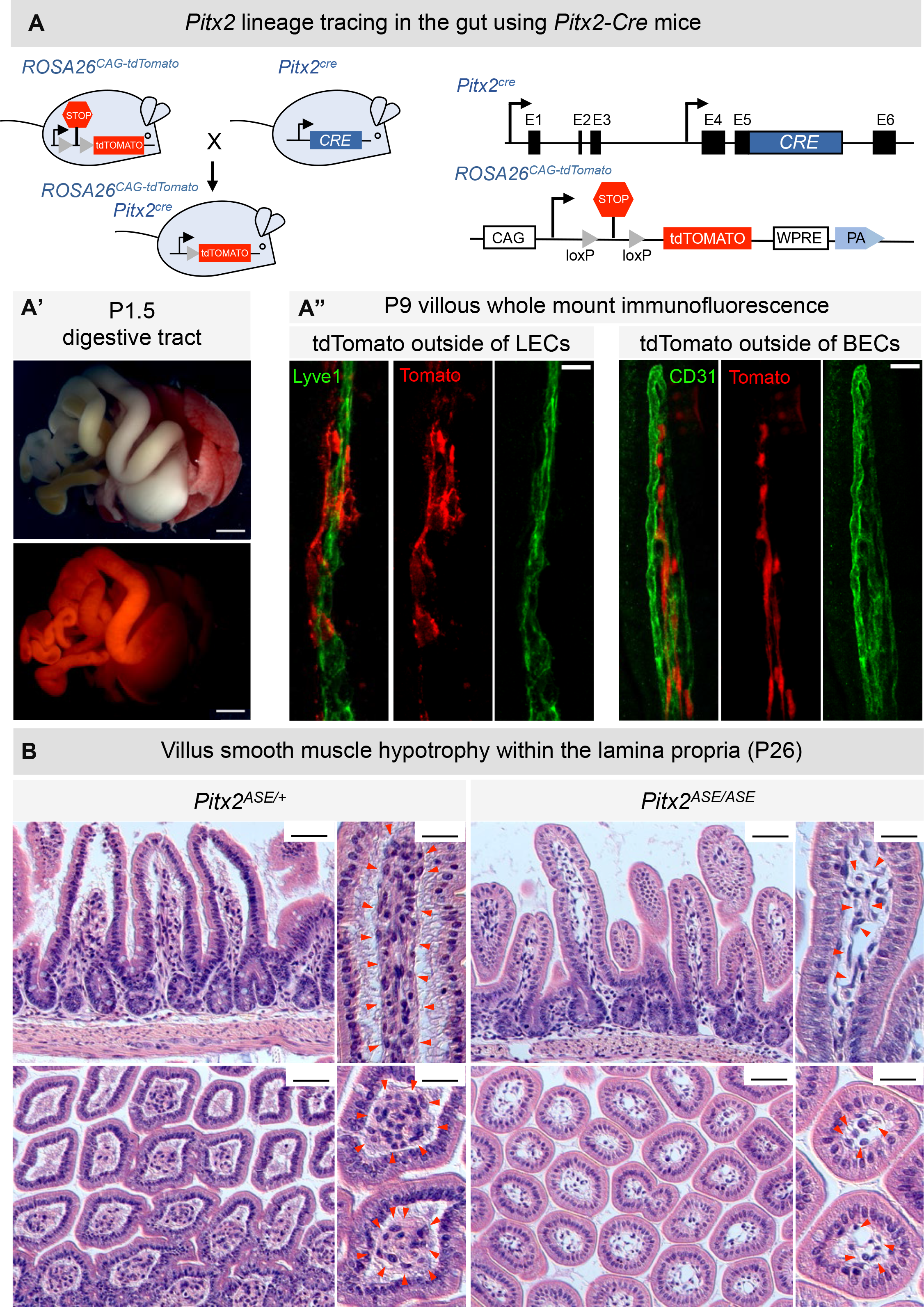
Villous SMCs are dependent on the *Pitx2* lineage (related to Figure 3) (A) Genetic background of *Pitx2^cre^* and *ROSA26^CAG-tdTomato^* mice used in lineage tracing. (A’) The tdTomato signal in a P1.5 *Pitx2^cre^*:: *ROSA26^CAG-tdTomatofl/fl^* mice is restricted to the digestive tract. Scale bar, 2000 μm. (A”) Representative images of villous whole mount IF in P9 *Pitx2^cre^*:: *ROSA26^CAG-tdTomato^* mice. <u>Left</u>: Green is lacteal (lyve-1) and red is tdTomato. <u>Right</u>: Green is blood capillary plexus (CD31) and red is tdTomato. Note that the tdTomato signal neither colocalized with lacteal lymphatic endothelial cells (LECs) nor blood endothelial cells (BECs). Scale bar, 20 μm. (B) H&E staining of paraffin-embedded sections reveals villus smooth muscle hypotrophy in *Pitx2^ASE/ASE^* proximal small intestine (P26). Note atrophy of eosinophilic stromal cells in *Pitx2^ASE/ASE^* mice marked by red dash lines and red arrowheads. Scale bar, 50 μm for low magnification images; 25 μm for high magnification images.

**Figure S4.**
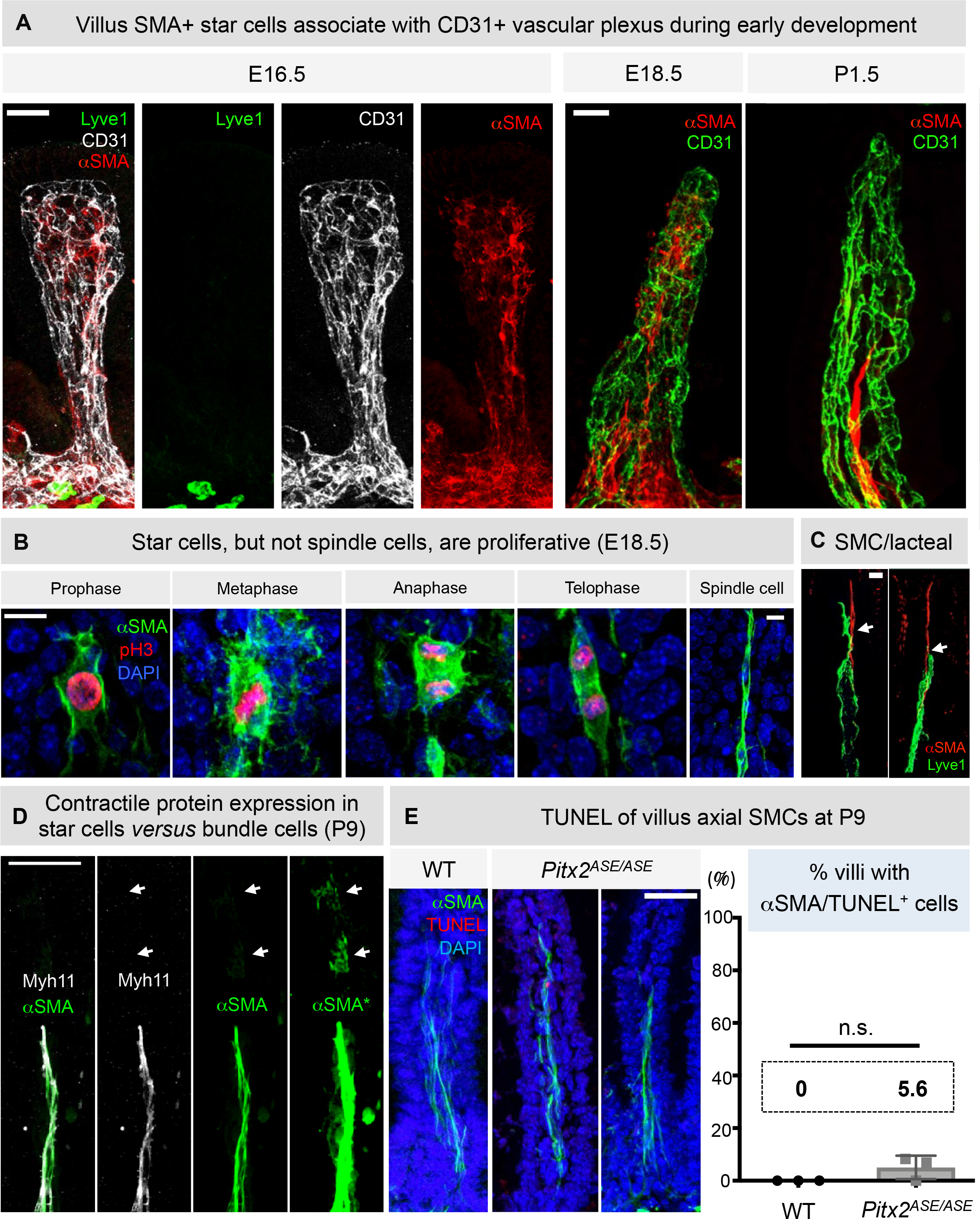
Lacteal-associated axial SMCs arise by *ASE*-dependent remodeling (related to Figure 4). (A) Villus SMA+ star cells associate with CD31+ vascular plexus during early development. <u>E16.5</u>: Representative images of whole mount WT villous processed for triple IF to demonstrate the close interaction of αSMA^+^ star cells (red) with CD31^+^ blood vascular plexus (white) in early lacteal development (lyve-1, green) and at <u>E18.5</u> (red is αSMA; green is CD31). <u>P1.5</u>: Note the αSMA^+^ star cells (red) seen at E16.5 and E18.5 are no longer seen at P1.5. Instead of star cells, αSMA^++^ spindle cells are observed resembling axial SMCs. Scale bar, 30 μm. (B) Star cells, but not spindle cells, are proliferative (E18.5). Representative images of whole mount WT villus; green is αSMA (green), red is phospho-histone 3 (H3P), and blue is DAPI. Note that pH3 signal colocalized only with star-like but not spindle-like SMCs. Additional panels are provided to show that star-like SMCs enter all four stages of mitosis (prophase, metaphase, anaphase, telophase). Scale bar, 10 μm. (C) Representative images of whole mount WT muscular-lacteal complexes at P1.5. Green is lacteal (lyve-1) and red is αSMA (smooth muscle). Note the closely intertwined axial SMCs with lacteal filopodia. Scale bar, 30 μm. (D) Representative images of villous whole mount IF at P9 demonstrating expression levels of contractile smooth muscle proteins αSMA (green) and myosin heavy chain 11 (Myh11, white). Note the intensity difference in contractile protein expression between star cells (white arrows) and axial SMC bundles. αSMA* marks higher exposure image needed to identify star cells. Scale bar, 40 μm. (E) *ASE* deletion does not result in increased cell death of axial SMCs. <u>Left</u>: Representative images of whole mount TUNEL assays on *Pitx2^ASE/ASE^* and WT villus at P9. Green is αSMA, red is TUNEL signal, and blue is DAPI. <u>Right</u>: % villi with TUNEL^+^/αSMA^+^ double-positive cells were quantified for both genotypes. Data displayed on the Y-axis are represented as mean ± SEM % of villi with TUNEL^+^/SMA^+^ double positive cells detected. WT = 0.00 ± 0.00 %, n = 3; *Pitx2^ASE/ASE^* = 5.56 ± 2.78 %, n = 3. Difference between WT and *Pitx2^ASE/ASE^* = 5.56 ± 2.78 %, P = 0.1161. Scale bar, 30 μm.

**Figure S5.**
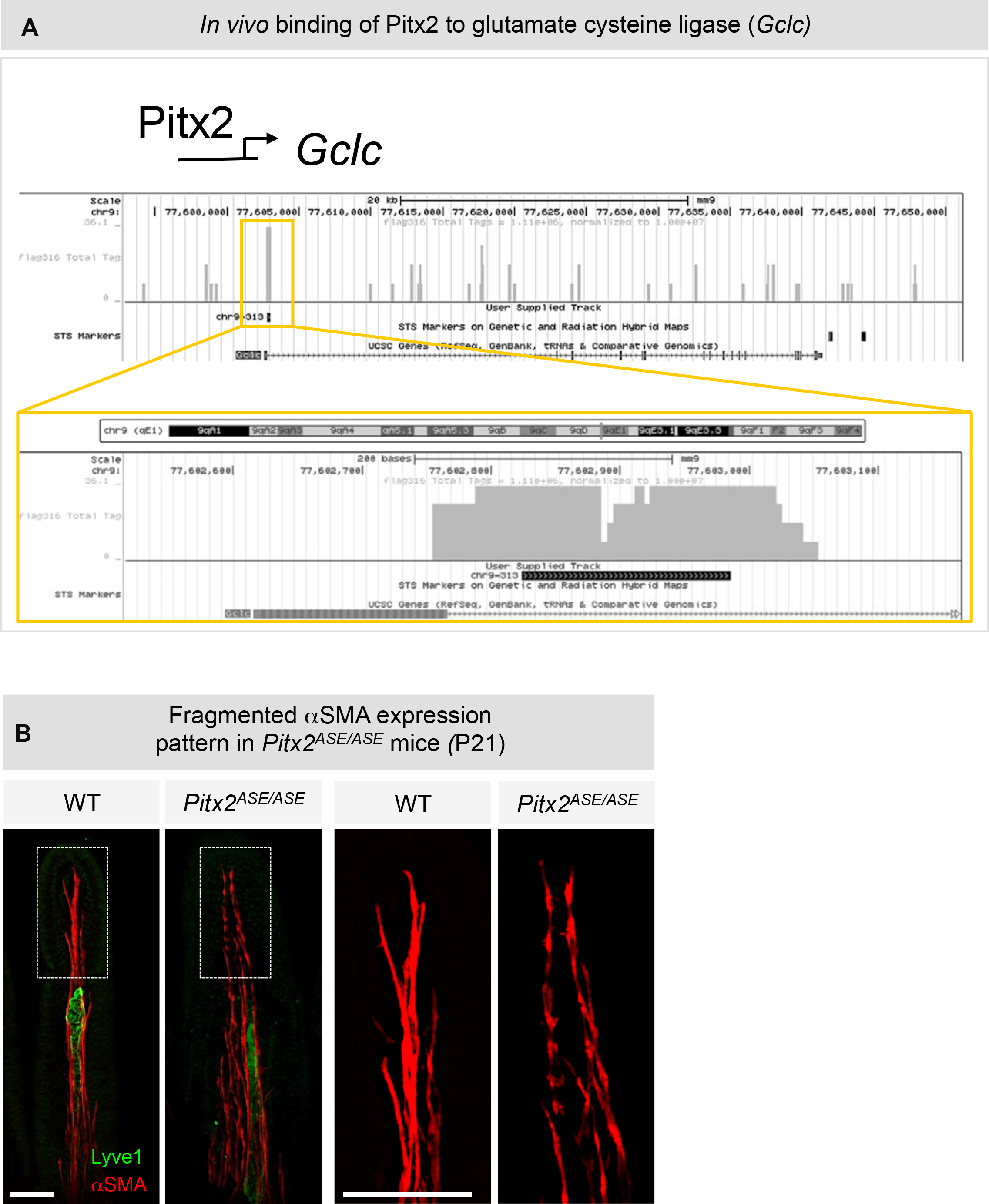
*ASE* deletion leads to redox imbalance and actin carbonylation in intestinal muscle (related to Figure 5) (A) Glutamate cysteine ligase *(Gclc)* was one of the 33 differentially expressed genes we have identified by RNA-seq of WT and *Pitx2^ASE/ASE^* intestines at P1.5. Pitx2 binds to the noncoding region of intron 1 (black box, Chr9-313) of *Gclc* as demonstrated by previous ChIP-seq. Scale bar, 20 kb (top) and 200 base pair (bottom). (B) Fragmented αSMA expression pattern in *Pitx2^ASE/ASE^* adult intestines (P21). Representative images of whole mount muscular-lacteal complexes of WT and *Pitx2^ASE/ASE^* littermates; green is lacteal (lyve-1); red is αSMA (smooth muscle). Note the continuous αSMA staining in WT *versus* fragmented pattern in *Pitx2^ASE/ASE^* axial SMCs. Dotted box represents area of higher magnification. Scale bar, 50 μm.

**Figure S6.**
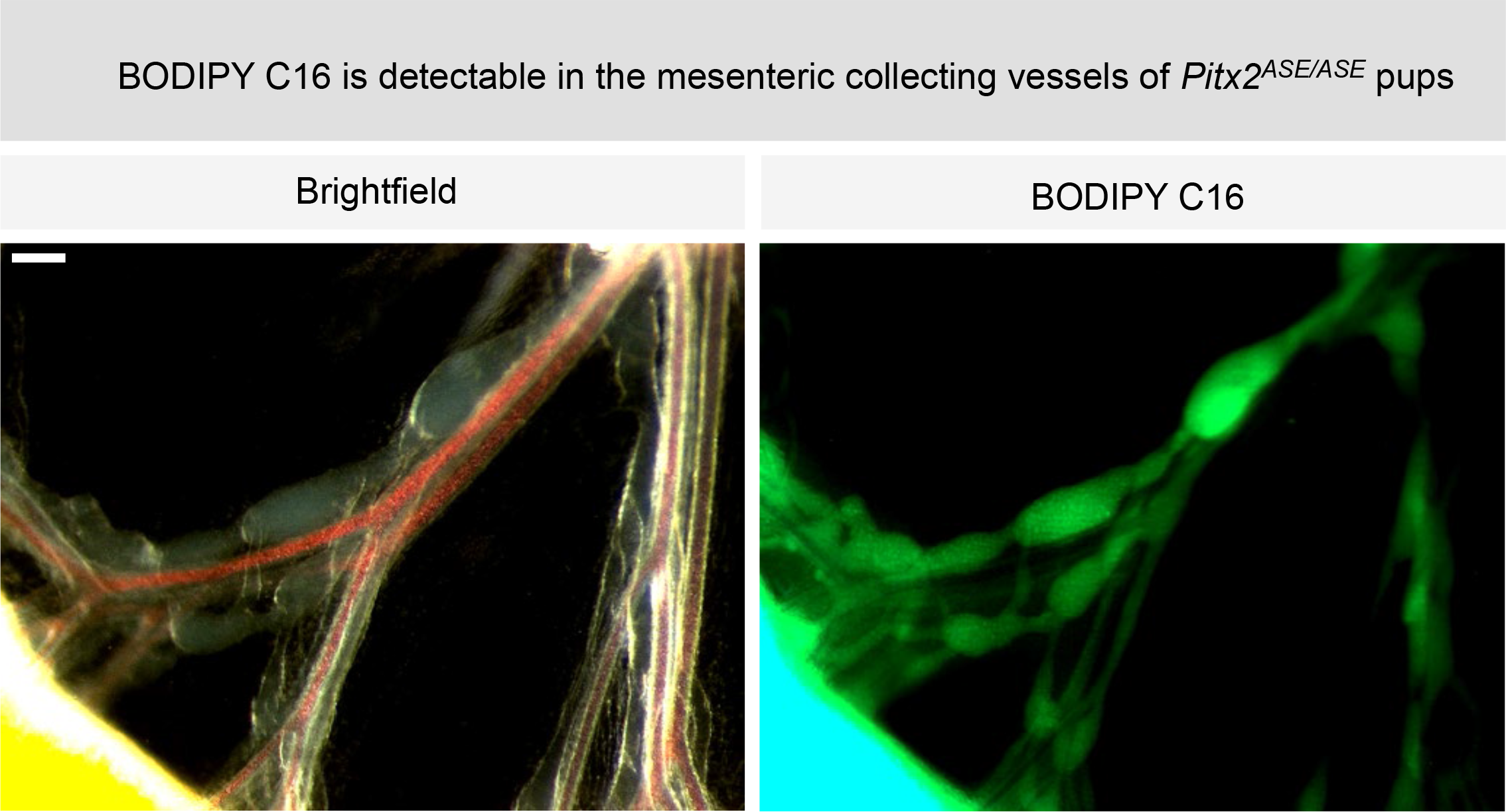
*ASE* is required for the normal route of dietary lipid transport (related to Figure 6) (A) Representative images of BODIPY™ FL C16 tracer entry into mesenteric collecting lymphatics in *Pitx2^ASE/ASE^* neonates. Images were taken 5 hours after feeding with the tracer. Signal in mesenteric collectors and lymphangions confirms the entry of dietary long-chain fatty acids into the lymphatic-dependent lipid transport pathway. Scale bar, 20 μm.

**Figure S7.**
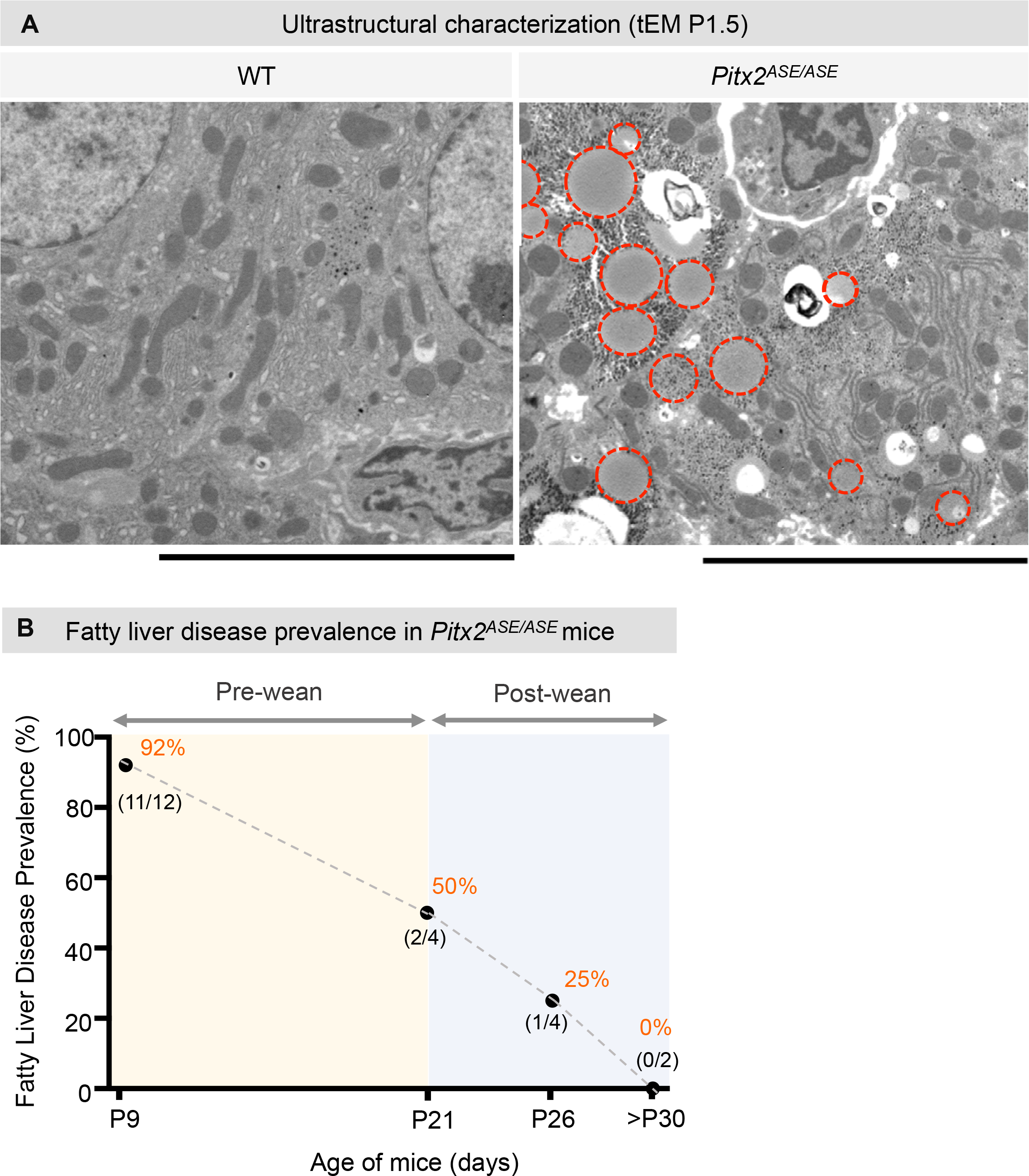
*Pitx2^ASE/ASE^* mice develop diet-induced fatty liver disease (related to Figure 7). (A) Representative images of tEM from WT (left) and *Pitx2^ASE/ASE^* (right) livers at P1.5. Note the accumulation of cytoplasmic lipid droplets (circled in red dash lines) in *Pitx2^ASE/ASE^* hepatocytes. Scale bar, 10 μm. (B) Graph demonstrating the % of *Pitx2^ASE/ASE^* mice with fatty liver disease lesions at pre-weaning (P9), weaning (P21), and post-weaning (P26-32). % prevalence of fatty liver disease is depicted in orange; black numbers depict positive cases of fatty liver disease over total cases examined. Data displayed on the Y-axis are represented as pooled fatty liver disease rate from n=12 (P9), n=4 (P21), n=4 (P26), n=1 (P31), and n=1 (P32) mice using histopathology examination.

## Notes

### Competing Interest Statement

The authors have declared no competing interest.

